# Protein structural transitions critically transform the network connectivity and viscoelasticity of RNA-binding protein condensates but RNA can prevent it

**DOI:** 10.1101/2022.03.30.486367

**Authors:** Andres R. Tejedor, Ignacio Sanchez-Burgos, Maria Estevez-Espinosa, Adiran Garaizar, Rosana Collepardo-Guevara, Jorge Ramirez, Jorge R. Espinosa

## Abstract

Biomolecular condensates, some of which are liquid-like during health, can age over time becoming gel-like pathological systems. One potential source of loss of liquid-like properties during ageing of RNA-binding protein condensates is the progressive formation of inter-protein β-sheets. To bridge microscopic understanding between accumulation of inter-protein β-sheets over time and the modulation of FUS and hnRNPA1 condensate viscoelasticity, we develop a multiscale simulation approach. Our method integrates atomistic simulations with sequence-dependent coarse-grained modelling of condensates that exhibit accumulation of inter-protein β-sheets over time. We reveal that inter-protein β-sheets notably increase condensate viscosity but does not transform the phase diagrams. Strikingly, the network of molecular connections within condensates is drastically altered, culminating in gelation when the network of strong β-sheets fully percolates. However, high concentrations of RNA decelerate the emergence of inter-protein β-sheets. Our study uncovers molecular and kinetic factors explaining how the accumulation of inter-protein β-sheets can trigger liquid-to-solid transitions in condensates, and suggests a potential mechanism to slow such transitions down.

## I. INTRODUCTION

Seen under a microscope, the eukaryotic cell would appear as an organised collection of billions of biomolecules exquisitely coordinated to carry out biological function and maintain cell structure [1, 2]. Compartmentalisation is a key feature in such coordination, and ensures that distinct regions in the cell are precisely enriched or depleted of specific molecules to fulfill their biological role [3, 4]. While the best-known form of these cellular compartments are membrane-bound organelles (e.g., the nucleus, the mitochondria [5, 6] or the Golgi apparatus [7]), the most widespread ones completely lack membranes [8, 9]. These membraneless compartments, known as biomolecular condensates, are formed by the process of liquid-liquid phase separation (LLPS), which is mainly driven by multivalent proteins and nucleic acids that can establish multiple homotypic or heterotypic interactions with cognate biomolecules (i.e., different proteins, RNA, or DNA) over their interactions with the surrounding media [10–14].

Phase separation is sensitive to thermodynamic conditions, which can be exploited by cells to react to environmental changes [15], such as temperature [16], salt gradients [17], pH [18, 19], or presence of malicious DNA in the cytosol (related to viral and microbial infections). This reliance on such a delicate equilibrium can cause misregulation of LLPS, promoting rigidification of liquid-like condensates into pathological solid aggregates [20–22]. Subtle changes in environmental conditions such as ionic salt concentration, pH, or decreased adenosine triphosphate (ATP) levels can give rise to decreased protein solubility [23]. In addition, post-translational modifications, such as phosphorylation [24] or specific protein mutations, can transform the binding affinity among species and critically alter the timescales of protein–protein interactions [25, 26]. For instance, mutations found in the fused in sarcoma (FUS) protein of amyotrophic lateral sclerosis (ALS) patients, significantly increase the rate and strength of its gelation [22, 27, 28]. Similarly, mutations in *α*-synuclein—a protein associated with Parkinson’s disease—can induce LLPS and subsequent ageing into gel-like droplets [29]. The formation of phase-separated nuclei of the Alzheimer-related τ-protein can also enhance the emergence rate of harmful amyloids [30]. However, condensate pathological solidification can also occur without the need of sequence mutations, post-translational modifications, external stimuli, or sensitive changes in the thermodynamic conditions [26, 31].

One of the proposed mechanisms to explain the liquid-to-solid mesoscale transformation of biomolecular condensates during ageing is the gradual accumulation of inter-protein structural transitions over time [32–37]. This is not surprising if one considers that the interaction landscape of proteins can be significantly transformed by structural transitions [33, 36–38]. Indeed, the low complexity domains (LCD) of various naturally occurring phase-separating proteins—including FUS [35], TAR DNA-binding Protein of 43 kDa (TDP-43) [39, 40], heterogeneous nuclear ribonucleoprotein A1 (hnRNPA1) [32–34], nucleoprotein of 98 kDa (NUP-98) [33, 41], and amyloid β (Aβ) NKGAII—contain short regions termed Low-complexity Aromatic-Rich Kinked Segments (LARKS), which are prone to forming inter-protein β-sheets in environments of high protein concentration [32, 42, 43]. These proteins form liquid-like condensates that can transition to hydrogels over time [44–46]. The inter-peptide *β*-sheet interactions are then thought to explain transient solidification of, otherwise, liquid-like condensates [33, 35–38, 47–49]. Importantly, hundreds of protein sequences capable of such structural transitions, and concomitant enhancement of inter-molecular binding strength, have been identified within the human genome [33].

Macroscopically, aged condensates can be unambiguously characterized by reduced fusion propensities and significantly longer recovery times [22, 29, 48, 50–53]. Techniques such as fluorescence recovery after photo-bleaching (FRAP) or green florescence protein (GFP) recovery have demonstrated that over time, even condensates that start displaying liquid-like behaviour can ‘age’ or ‘mature’ (i.e., change their material properties), transitioning into gels or soft glasses [20, 21, 25]. Notably, particle tracking microrheology techniques have been also successfully employed to evaluate the mean squared displacement (MSD) of marked beads inside droplets, and then, via that MSD and the Stokes-Einstein relation, the viscosity of the condensates can be inferred [25, 54–57]. Moreover, the progressive dynamical arrest of proteins has been also observed *in vitro* for protein condensates containing marked prion-like domains (PLDs) enriched in LARKS [3, 19, 21, 22, 33, 35, 36, 45, 58–61]. Nevertheless, characterising the microscopic origin and molecular mechanisms by which condensates age over time, still remains extremely challenging [62–64].

In that respect, computational approaches may provide insightful guidance on the thermodynamic and molecular driving forces underlying condensate pathological ageing [65, 66]. From atomistic simulations [36, 67–70] to coarse-grained models [71–79] including lattice-based simulations [80–82] and mean-field theory [83–85], computational science has significantly contributed to understanding the role of RNA in regulating the dynamics of multivalent phase-separared droplets [86–88], the impact of strong-binding in condensate rigidification [37, 38] or the formation of kinetically-arrested multiphase condensates from single-component droplets [36]. Nevertheless, further insights on the different possible causes behind condensate ageing—e.g., molecular inter-protein binding events, amino acid sequence mutations, or relevant variations on the applied thermodynamic conditions—are urgently required. In that sense, proposing effective strategies to liquefy biomolecular condensates has become a key area of research to prevent the proliferation of neurodegenerative disorders [9, 31, 89–91]—such as amyotrophic lateral sclerosis (ALS) [92], Parkinson’s [29], Alzheimer’s [30] or frontotemporal dementia (FTD)—as well as certain types of cancers [93] or diabetes [94] associated to the progressive formation of solid-like aggregates.

In this work, we develop a multiscale computational approach, integrating atomistic simulations and residuere-solution coarse-grained models to shed light on the thermodynamic and kinetic factors that explain ageing of biomolecular condensates via gradual accumulation of β-sheet content in the presence and absence of RNA. We compare the behaviour of FUS condensates *versus* hnRNPA1 condensates (A1-A isoform, or A-LCD-hnRNPA1) because of their relevance to the formation of stress granules [21, 32, 95], the so-called ‘crucible for ALS pathogenesis [96]’. We first use atomistic simulations to quantify the change in the binding free energies of proteins as a result of inter-protein β-sheet assembly. We then investigate the nonequilibrium process of condensate ageing using a residue-resolution coarse-grained model that considers our atomistic results. To model ageing, we develop a dynamical algorithm that models the time-dependent formation of inter-protein β-sheets inside condensates. We consider that condensate ageing is a nonequilibrium process where there is a gradual increase in the imbalance of inter-molecular forces (i.e., these are nonconservative) over time. We find that while accumulation of long-lived inter-protein β-sheets only moderately increases the density of condensates, its impact on their viscosity is compelling, especially at low temperatures. Strikingly, recruitment of a high concentration of RNA into the condensates hampers the size and concentration of the inter-protein β-sheet nuclei. The disrupting effect of RNA–RNA repulsive interactions on the condensate liquid-network connectivity [97], in addition to the variation of the molecular contact amalgam sustaining LLPS, effectively precludes the spontaneous emergence of high-density protein regions prone to assemble into strongly binding domains through inter-protein *β*-sheet transitions. Such reduction in inter-protein disorder-to-order transitions considerably improves the liquid-like behaviour of the condensates by decreasing their viscosity and density. Our study therefore contributes to rationalizing the microscopic origin by which naturally occurring proteins—capable of exhibiting structural transitions [33]—may progressively drive biomolecular condensates into kinetically-arrested aggregates, and suggests an effective mechanism by which condensates may actively avoid undergoing pathological liquid-to-solid transitions.

## II. RESULTS

### A. Inter-peptide β-sheet clusters critically enhance protein binding

We begin by estimating the changes to the free energy of binding among proteins when they transition from disordered to inter-protein β-sheets. We focus on the behaviour of hnRNPA1 and FUS, as these are two naturally occurring phase-separating RNA-binding proteins that form condensates which are prone to undergo ageing. For this, we perform atomistic Umbrella Sampling Molecular Dynamics (MD) simulations [98] of systems containing four identical hnRNPA1 interacting LARKS peptides (_58_GYNGFG_63_ [33]) in explicit solvent and ions at room temperature and physiological salt concentration using the all-atom a99SB-disp force field [99]. For FUS, we compare with our results for the three different LARKS found in the LCD (i.e., _37_SYSGYS_42_, _54_SYSSYGQS_61_, and _77_STGGYG_82_ [35]) using both a99SB-disp [99] and CHARMM36m force fields [100], reported in Ref. [36]. For each case, we calculate the free energy cost of dissociating one single segment from a cluster containing four identical peptides under two distinct scenarios: (1) when the four peptides are fully disordered; and (2) when the four peptides instead form a cross-β-sheet motif resolved crystallographically (e.g., PDB code: 6BXX for hnRNPA1 [33]). We compute the Potential of Mean Force (PMF) as a function of the centre-of-mass (COM) distance between one single peptide—which we gradually dissociate from the other segments—and the other three peptides (simulation details are described in the Supplementary Methods). In the initial scenario in which LARKS are treated as fully disordered segments, we allow peptides to freely sample their conformational space (only fixing the position, in the appropriate direction, of the closest atom to the peptide COM of the structured four-peptide motif; see Supplementary Material for further details). In the second scenario in which we quantify the interactions among structured LARKS, we constrain the peptides to retain their crystal β-sheet structure as described in Refs. [17, 36–38, 101].

Consistent with the liquid-like behaviour for A-LCD-hnRNPA1 condensates [51, 102, 103], our simulations reveal that the binding strength among fully disordered hnRNPA1 LARKS is sufficiently weak (i.e., ~1 k_*B*_T per residue) for their interaction to be transient (Fig. 1a, green curve). However, when such LARKS assemble into ordered inter-protein β-sheet structures, their binding strength increases by 400% (i.e., >4 k_*B*_T per residue, Fig. 1a; yellow curve). FUS was previously found to exhibit the same behaviour, with the strength of inter-peptide interaction increasing significantly upon inter-peptide β-sheet formation (i.e., from 1–2 k_*B*_T to 4.5–5.6k_*B*_T per residue) [36]. We note that the exact magnitude of this increase might be slightly overestimated due to the required constraints to enforce the stability of the β-sheet structures. Nevertheless, considering only the impact of imposing positional restraints in PMF simulations is not sufficient to account for the observed increase in binding strength [36–38]. Both the specific inter-protein secondary structure and the amino acid composition are crucial parameters to enable strengthened protein binding via inter-peptide β-sheet domains as previously demonstrated in Refs. [36, 37]. Importantly, the relatively high interaction strengths among structured LARKS that we obtain here for hnRNPA1 (~25k_*B*_T for the whole peptide), and previously for FUS (up to ~40k_*B*_T per peptide), are consistent with the formation of reversible hydrogels that can be easily dissolved with heat, as found experimentally [33, 35, 39]. Thermostable amyloid fibrils, such as those formed by the Aβ1–42, are expected to be stabilized by considerably larger binding energies, e.g. of the order of 50-80 k_*B*_T [33, 38, 104–106]. Overall, our atomistic results highlight how a critical enhancement of interactions among RNA-binding proteins in phase-separated condensates can occur in absence of chemical modifications or variations in the thermodynamic conditions (i.e., temperature, pH or salt gradients), and be driven by the formation of inter-peptide LARKS β-sheets.

**Figure 1:**
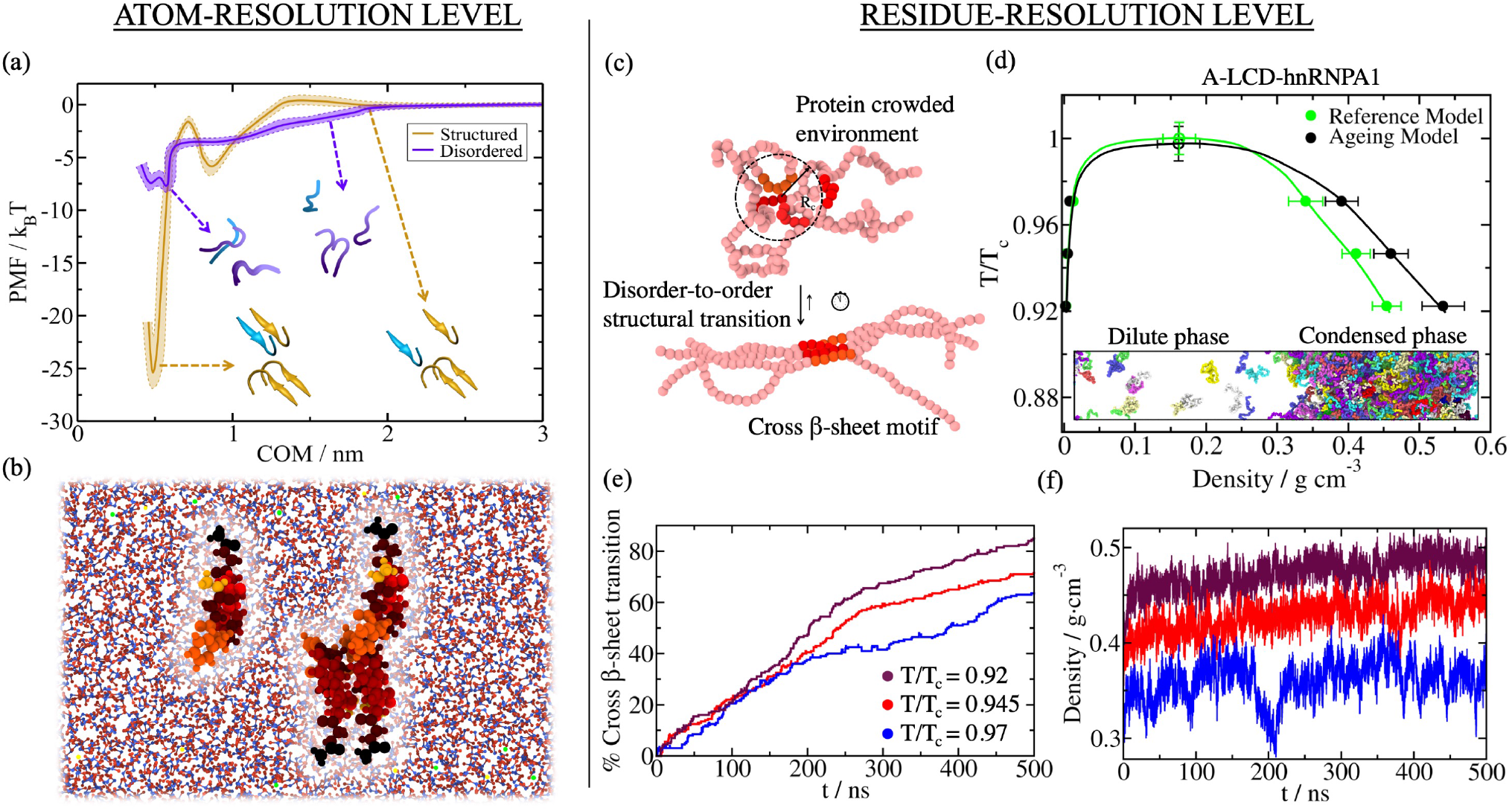
Structural transitions leading to inter-peptide β-sheet motifs dramatically increase protein binding and promotes droplet densification over time. (a) Atomistic Potential of Mean Force (PMF) dissociation curve of a 6-amino acid segment (PDB code: 6BXX) found in the A-LCD-hnRNPA1 sequence from a β-sheet structure formed by 4 peptides (of the same sequence) as a function of the center of mass distance (COM) using the a99SB-disp force field [99]. PMF simulations are conducted at room conditions and physiological salt concentration. Yellow curve represents the interaction strength among peptides with a well-defined folded structure, kinked β-sheet structure, while the purple curve depicts the interaction strength among the same segments but when they are fully disordered. Statistical uncertainty is depicted by colour bands. A representation of the four peptides, both ordered (yellow) and disordered (purple) including the dissociating peptide in blue, is also included in the inset. (b) Snapshot of an all-atom PMF simulation in which the structured peptide is pulled from the inter-protein β-sheet motif. The distinct residues within the peptides are highlighted by different colours while water is depicted in blue (O) and red (H) and NaCl ions by green (Na^+^) and yellow (Cl^-^) spheres. (c) Representation of the dynamical algorithm coupled to the residue-resolution model to introduce disorder-to-order transitions according to the protein local environment. When four LARKS segments meet within a given cut-off distance, LARKS binding is strengthened according to the PMF binding free energy difference computed in Panel (a). The bending penalty between residues composing LARKS motifs is also enhanced to account for the higher rigidity of structured β-sheet aggregates [33, 35]. (d) Phase diagram of A-LCD-hnRNPA1 in the T-*ρ* plane for protein condensates with disorder-to-order transitions and subsequent strengthening of inter-molecular protein binding (black symbols; dynamical ageing model), and for the reference model (HPS-Cation-π, Ref. [107, 108]) where the interaction strength among LARKS is always considered fully disordered (green symbols). Statistical errors are obtained by bootstrapping results from n = 3 independent simulations. A Direct Coexistence simulation snapshot is included in the inset where different protein replicas are depicted by different colours. (e) Number of inter-peptide (cross) β-sheet transitions as a function of time found in phase-separated condensates at different temperatures (see legend; temperatures are normalized by the critical temperature of A-LCD-hnRNPA1; T_*c*_). (f) Time-evolution of the condensate density for different temperatures as indicated in the legend of Panel (e).

### B. Time-dependent modulation of protein phase diagrams during ageing

The high protein concentrations found inside condensates are expected to facilitate the interaction between multiple LARKS, and as a result, encourage the formation of inter-protein β-sheets. Motivated by this, we explore how our atomistic observations for a few interacting LARKS peptides (i.e., strengthening of interactions due to inter-peptide β-sheet ladders assembly) would impact the behaviour of condensates containing high RNA-binding protein concentrations. For this, we develop an innovative multiscale simulation approach that integrates our atomistic LARKS–LARKS binding free energies (from Fig. 1 and Ref. [36]), a residue-resolution coarse-grained protein model [107, 108, 111, 112], and a dynamical algorithm that we develop here to describe the nonequilibrium process of condensate ageing due to inter-peptide β-sheet formation. Coupled to the residue-resolution model [107, 108, 111], our dynamic algorithm approximates the process of condensate ageing by considering the atomistic implications (i.e., non-conservative strengthening of inter-protein binding, local protein rigidification, and changes in the inter-molecular organization) of the gradual and irreversible accumulation of inter-protein β-sheet structures in a time-dependent manner, and as a function of the local protein density within phase-separated biomolecular condensates (Fig. 1c).

In more detail, we describe the potential energy of interacting coarse-grained proteins (A-LCD-hnRNPA1 and FUS) within a condensate prior to ageing with the HPS-Cation-π [107] re-parameterization of the HPS model [108]. Recently, we showed that the HPS-Cation-π parametrization qualitatively reproduces the relative propensity of numerous RNA-binding proteins (including FUS and A-LCD-hnRNPA1) to phase separate at physiological conditions [87], as well as their RNA-concentration-dependent re-entrant behaviour [113–116]. To enable condensate ageing, we introduce our dynamical algorithm that triggers transitions from disordered peptides to inter-protein β-sheets within selected LARKS (which now are not isolated peptides, but part of the whole A-LCD-hnRNPA1 or FUS protein sequence) when the central C_*α*_ bead of a LARKS is in close contact (within a cut-off distance of ~8Å) with three other LARKS of neighbouring proteins [33, 35, 39]. Every 100 simulation timesteps, our dynamical algorithm evaluates whether the conditions around each fully disordered LARKS are favorable for undergoing an ‘effective’ disorder-to-order cross-*β*-sheet transition. An ‘effective’ structural transition is defined as one that is enforced and recapitulated in our algorithm by enhancing the interaction strength of the four involved LARKS–LARKS pairs by a given factor according to our atomistic PMF simulations in Fig. 1a and Ref. [36]. In Supplementary Figure 1, we demonstrate how a PMF dissociation curve of the two sets of parameters in our coarse-grained model (Reference model: mimicking interactions among disordered protein regions vs. Ageing model: describing interactions among protein regions forming inter-protein β-sheets) recapitulates the atomistic binding free energies estimated for hnRNPA1 (58GYNGFG63 LARKS sequence; Fig. 1a). Therefore, by employing the coarse-grained model, we can perform Direct Coexistence simulations [117, 118] to estimate and compare the phase diagrams of A-LCD-hnRNPA1 and FUS, prior and post-ageing using tens to hundreds of protein replicas (Figs. 1d and 2a). Further details on the dynamical algorithm, the local order parameter driving structural transitions, PMF coarsegrained calculations and the structured interaction parameters of the coarse-grained model are provided in the Supplementary Methods.

**Figure 2:**
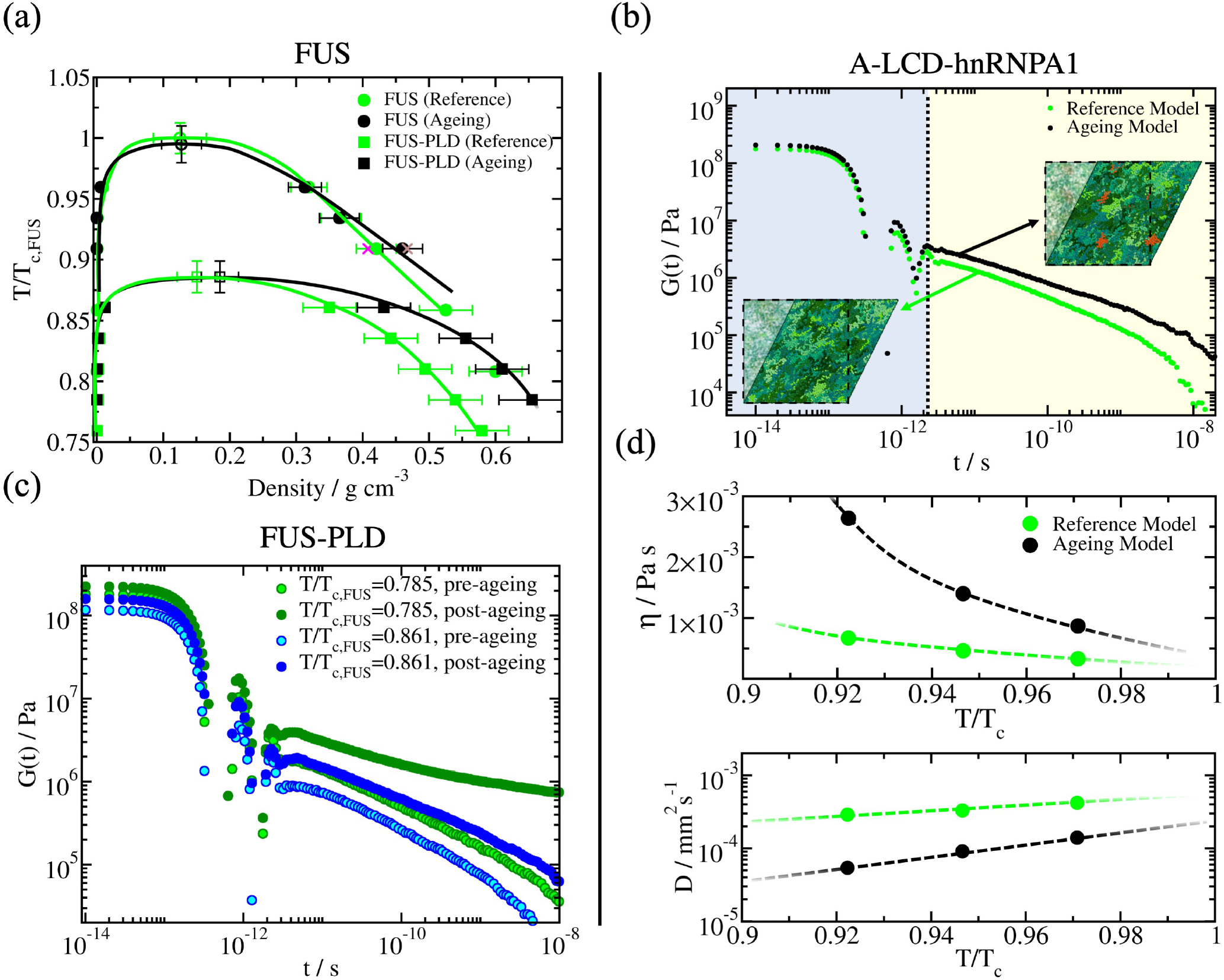
Protein structural transitions severely impact the viscoelastic behaviour of FUS and A-LCD-hnRNPA1 condensates. (a) Phase diagram in the T-*ρ* plane for full-FUS (circles) and FUS-PLD (squares) sequences before disorder-to-β-sheet transitions take place (green symbols), and after condensates become kinetically-arrested (black symbols; dynamical algorithm). Filled symbols represent the coexistence densities obtained via Direct Coexistence simulations [109], while empty symbols depict the estimated critical points by means of the law of rectilinear diameters and critical exponents [110]. Purple and brown crosses depict coexistence densities for the full-FUS reference model and ageing model respectively using system sizes two times larger (i.e., 96 protein replicas). Temperature has been normalized by the critical temperature of full-FUS (T_*c,FUS*_) for the reference model [107, 108]. Statistical errors are obtained by bootstrapping results from n = 3 independent simulations. (b) Shear stress relaxation modulus *G*(*t*) of the A-LCD-hnRNPA1 bulk condensed phase at T=0.92T_*c*_ for the reference model (purple curve; HPS-Cation-π model [107, 108]), and for protein condensates in which local strengthening of LARKS protein binding due to structural transitions is accounted (black curve; dynamical ageing model). Snapshots illustrating a shear stress relaxation computational experiment over A-LCD-hnRNPA1 condensates are included: Bottom, for a liquid-like condensate (reference model), and Top, for an aged condensate. Structured inter-peptide β-sheet motifs are depicted in red, and intrinsically disordered regions in green. (c) Shear stress relaxation modulus *G*(*t*) of FUS-PLD bulk condensates at two different temperatures as indicated in the legend. Light-coloured circles account for condensates before exhibiting binding strengthening due to structural transitions (reference model), while dark-coloured circles represent *G*(*t*) for condensates upon ageing (i.e., when the rate of structural transitions has reached a plateau). (d) Top: Viscosity (*η*) as a function of temperature (renormalized by the critical temperature T_*c*_) for A-LCD-hnRNPA1 condensates before the emergence of enhanced binding due to local structural transitions (green symbols; reference model), and after the formation of inter-protein β-sheet fibrils within the condensates (black symbols; dynamical model). Bottom: Protein diffusion coefficient within the bulk condensed phase before (green symbols) and after the formation of inter-peptide β-sheet motifs within the condensates (black symbols).

When we investigate the impact of ageing on the phase diagrams (in the temperature–density plane) of A-LCD-hnRNPA1, full FUS, and the PLD of FUS, our Direct Coexistence simulations consistently predict that, in all cases, the critical parameters of the different proteins are not affected by ageing (Fig. 1d and Fig. 2a). In the Direct Coexistence method, the two coexisting phases are simulated by preparing periodically extended slabs of the two phases (the condensed and the diluted phase) in the same simulation box. Once the system reaches equilibrium (or a steady state in the case of non-equilibrium systems), density profiles along the long axis of the box can be extracted to compute the density of the two coexisting phases. We note that when ageing is driven by inter-protein fibrillization, like in our simulations, conservation of critical parameters during ageing is not entirely surprising: for systems exhibiting upper-critical solution temperatures, the crucial structural transitions that drive ageing are gradually disfavoured as we approach critical conditions (Fig. 1e) because of the progressively decreasing protein densities found in the condensate at higher temperatures—shown in Fig. 1f. While the critical parameters are conserved, the faster accumulation of inter-protein β-sheets (where % Cross-β-sheet transition refers to the number of emerged transitions over the total number of LARKS in our system, i.e. the number of protein replicas times the number of LARKS per protein replica) at decreasing temperatures (Fig. 1e) results in increasingly divergent material properties of the aged versus pre-aged condensates. That is, aged condensates (Fig. 1d and Fig. 2a; black symbols) are denser than their pre-aged counterparts (Fig. 1d and Fig. 2a; green symbols), and this difference increases gradually as the temperature decreases. In the case of FUS, the increase in density upon ageing is modest for the full protein but significant for the FUS-PLD—we attribute this to three LARKS being contained within the short FUS-PLD of 163 residues versus the longer full FUS protein (526 amino acids in total). Independently of temperature, in all cases, we observe that the number of inter-protein β-sheets in the condensate increases over time, and consequently also the density of the aged condensates (Figs. 1f and 2a). These results are consistent with recent experimental observations of FUS condensates, where an increase in the β-sheet content induces a rise in droplet density upon thermal-annealing induced ageing [43].

### C. Emergence of inter-protein β-sheets during ageing increases droplet viscosity

Our observations that the extent of ageing and its effects in modulating the density of condensates is amplified at lower temperatures, motivated us to next investigate the impact of ageing in the viscoelastic properties of condensates as a function of temperature. The time-dependent mechanical response of a viscoelastic material when it is subjected to a small shear deformation can be described by the relaxation modulus *G*(*t*) [119]. This relaxation modulus can be determined by computing the auto-correlation of any of the off-diagonal components of the pressure tensor. If the system is isotropic, a more accurate expression of *G*(*t*) can be obtained by using the six independent components of the pressure tensor, as shown in Ref. [120] (see Supplementary Methods for further details on this calculation). The zero-shear-rate viscosity of the system can be computed by integrating in time the stress relaxation modulus. The direct evaluation of *G*(*t*) from our simulations provides critical information not only on how the material properties of condensates change during ageing, but also on how such changes are dictated by different relaxation mechanisms of the proteins that compose them (Fig. 2b). At short timescales (light blue region), the stress relaxation modulus mostly depends on the formation and breakage of short-range interactions and on intra-molecular reorganisation (i.e., internal protein conformational fluctuations, such as bond or angle relaxation modes). At long timescales (beige region), the stress relaxation modulus is mainly dominated by inter-molecular forces, long-range conformational changes (i.e. protein folding/unfolding events), and protein diffusion within the crowded liquid-like environment of the condensate.

Looking at the time-dependent behaviour of the stress modulus for A-LCD-hnRNPA1 condensates at a sub-critical temperature (i.e., T/T_*c*_=0.92), we observe a consistent decay of *G*(*t*) over time for both aged and pre-aged condensates, indicative of the liquid-like character of both condensates (Fig. 2b). Despite this apparent similarity, ageing of A-LCD-hnRNPA1 condensates slows down the rate of decay of *G*(*t*), which signals a higher viscosity for aged condensates due to the strengthening of inter-molecular forces as inter-protein β-sheets accumulate (Fig. 2b; black curve). Such a higher viscosity of aged condensates is increasingly accentuated as the temperature decreases (Fig. 2d (Top panel); i.e., below T/T_*c*_=0.9) and the structural transitions are favoured (Fig. 1e). We note that the coarse-grained nature of our implicit-solvent model [107, 108, 111] can significantly underestimate the relaxation timescale of the proteins, and hence, droplet viscosity [87]; however, the observed trends and relative differences in viscoelastic properties among pre-aged and aged condensates are expected to hold despite the artificially faster dynamics of our residue-resolution simulations.

In the case of pre-aged FUS-PLD condensates, the stress relaxation modulus decays continuously over time, demonstrating, as in the case of A-LCD-hnRNPA1, liquid-like behaviour prior to ageing (Fig. 2c). While the decay of G(t) over time emerges at different sub-critical temperatures (T/T_*c,FUS*_=0.785 and T/T_*c,FUS*_=0.861, where the critical temperature of FUS-PLD corresponds to T_*c,FUS–PLD*_ ≈0.875 T_*c,FUS*_), it exhibits considerably shorter relaxation times at the higher temperature. Moving now towards aged FUS-PLD condensates, we observe that, irrespective of temperature, ageing increases significantly the values of the shear stress relaxation modulus; hence, suggesting a much higher viscosity for FUS-PLD aged condensates than their pre-aged counterparts. However, when looking more closely at the time-dependent behaviour of *G*(*t*), we observe significantly different profiles at varying temperatures. At high temperatures (e.g., T/T_*c,FUS*_=0.861), the continuous decay of *G*(*t*) for aged FUS-PLD condensates is consistent with that of a liquid (Fig. 2c; light blue circles). Yet, at lower temperatures (e.g., T/T_*c,FUS*_=0.785), *G*(*t*) falls into a persistent plateau with no hints of decaying at comparable timescales, and yielding infinite viscosity values (and non-diffusive behaviour) characteristic of a gel-like state as recently reported in Ref. [25] for FUS condensates.

When assessing the mobility of A-LCD-hnRNPA1 proteins inside pre-aged condensates, our simulations reveal a constant liquid-like behaviour along the entire range of sub-critical temperatures we study, consistent with the behaviour of *G*(*t*) (Fig. 2d; Top panel). In contrast, and also in agreement with our viscosity measurements, protein mobility within aged condensates is severely limited (Fig. 2d; Bottom panel). The deceleration of protein diffusion in aged condensates becomes more pronounced as the temperature decreases, as expected from the faster accumulation of inter-protein β-sheets at lower temperatures (Fig. 1e). In FUS and FUS-PLD aged condensates, viscosity and diffusion coefficients cannot be reliably evaluated due to the higher number of LARKS within the sequence (3 domains) compared to A-LCD-hnRNPA1 (1 domain) impeding the full relaxation of *G*(*t*) over time (i.e., non-ergodic gel-like behaviour is observed as shown in Fig. 2c; dark green circles). In experiments, a decrease of protein mobility over time during ageing is reflected in decreased diffusion coefficients, higher condensate viscosities [29, 121] and lower or incomplete recovery after photobleaching [29, 31, 50–52, 122, 123]. Moreover, in line with our results, slowdowns in protein mobility have been observed experimentally for aged hnRNPA1 and FUS phase-separated droplets [25, 32–34]. Remarkably, in some of those studies, the reported deceleration in protein mobility has been associated with a significant increase of β-sheet content within the condensates [32, 43], which emphasizes the strong interplay between protein mobility and inter-protein β-sheet content.

### D. Liquid-like and gel-like aged condensates present drastic differences in their networks of inter-molecular connections

The striking temperature-dependent changes to the behaviour of the stress relaxation modulus for FUS-PLD suggest a potential transformation of the condensate percolating network of inter-molecular binding when the condensate transitions from a liquid into a gel (i.e., at T/T_*c,FUS*_=0.785; Fig. 2c). To characterise the structure and topology of the condensate inter-molecular network, we develop a modification to the primitive path analysis algorithm originally proposed to reveal the underlying network structure of polymer melts [124, 125]. In our method, we consider that β-sheet LARKS–LARKS bonds are fixed in space, the intra-molecular excluded volume is set to zero, and the bond interaction is modified to have an equilibrium bond length of 0 nm. This algorithm minimises the contour length of the protein strands that connect the different LARKS regions, while preserving the topology of the underlying network, and allows for visualisation of the network connectivity generated by the inter-protein β-sheet clusters (Fig. 3c, please see Supplementary Methods and Supplementary movies 1 and 2 for further details). Furthermore, to better observe the extension of the network connectivity beyond the periodic boundary conditions of the simulation box, we replicate the system in all directions of space. At the end of the minimization, we can observe the network of elastically active protein strands that contributes to the formation of a rubbery plateau in *G*(*t*) (as shown in Fig. 2c; dark green circles). If this network percolates, the relaxation modulus will show a clear plateau, whereas if the proteins form disconnected clusters, *G*(*t*) will decay to zero—although it will still exhibit higher viscosity compared to the non-aged condensates sustained by short-lived inter-protein bonds.

**Figure 3:**
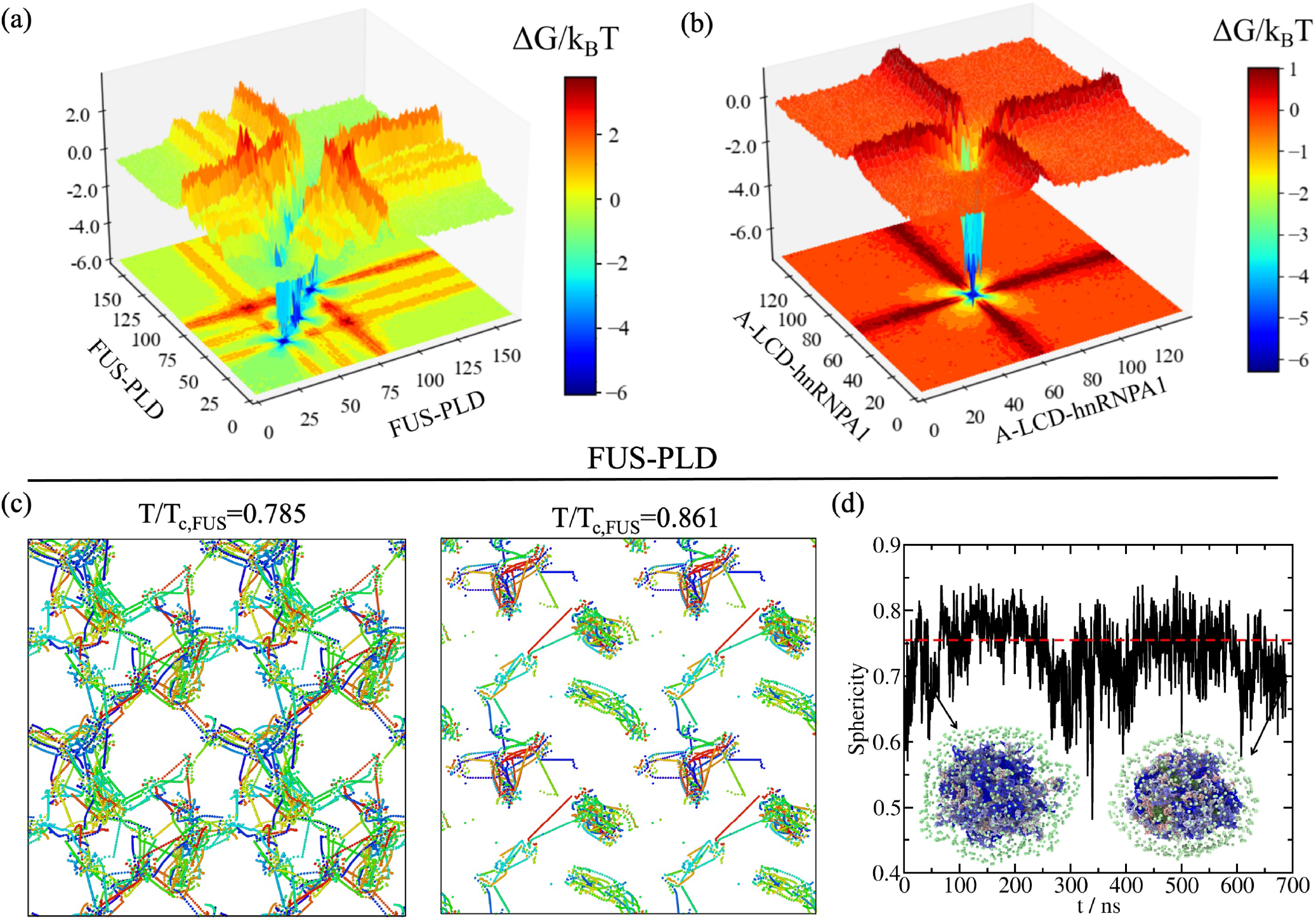
Imbalanced protein binding can drive condensate ageing but not droplet reshaping. Landscape of the protein contact free energy variation upon condensate ageing for bulk FUS-PLD condensates at T/T_*c,FUS*_=0.785 (a) and A-LCD-hnRNPA1 condensates T=0.97T_*c*_ (b). ΔG/k_*B*_T is obtained from the residue contact probability ratio between aged condensates and liquid-like condensates before ageing. Colour map projections of the free energy landscape in 2-dimensions are also included. (c) Network connectivity of aged FUS-PLD condensates at T/T_*c,FUS*_=0.785 (Left) and T/T_*c,FUS*_=0.861 (Right) computed using a primitive path analysis. (d) Evolution of droplet sphericity during condensate ageing at T/T_*c,FUS*_=0.81. The percentage of transitioned fully-disorder LARKS into structured inter-peptide β-sheet motifs was ~75% along the trajectory. The horizontal red dashed line represents the average sphericity for a FUS-PLD condensate in a liquid-like state (reference model). Two representative configurations of the condensate surrounded by the surface map (light green) are provided with LARKS residues belonging to inter-peptide β-sheet clusters depicted in dark green and fully disorder residues belonging to distinct protein replicas depicted by a blue-to-grey colour range. Details on the sphericity parameter evaluated through a Solvent Available Surface Area (SASA) analysis as well as of the primitive path analysis and the protein contact free energy calculations are provided in the Supplementary Methods.

Our results reveal a remarkable transformation of the molecular connectivity of the aged condensate at lower temperatures that is required to enable its gelation. At the lowest temperature (T/T_*c,FUS*_=0.785), where the viscoelastic properties of aged FUS-PLD condensates are consistent with a gel, we observe a network of strong inter-protein β-sheet contacts that completely percolates through the aged FUS-PLD condensate. Such a fully connected network of strong inter-molecular bonds inhibits the relaxation of molecules within the condensate at long timescales, leading to its gel-like properties, such as a stress relaxation modulus that reaches a plateau (dark green circles; Fig. 2c). At higher temperatures (T/T_*c,FUS*_=0.861) where aged FUS-PLD condensates remain liquid-like, the network of inter-protein β-sheet contacts in the aged FUS-PLD condensate presents only isolated gel-like structures; it is precisely the lack of full percolation in strong β-sheet contacts what allows the aged condensate to relax as a whole and behave as a high viscosity liquid (dark blue circles; Fig. 2c).

Importantly, a fundamental requirement for proteins to exhibit the type of gelation upon ageing that we describe here for FUS-PLD droplets is to have at least three separate LARKS segments. This is because the described gelation emerges only when the strong β-sheet LARKS–LARKS bonds can form a state of full connection; in other words, at least three anchoring points per molecule are necessary for a system to completely gelate [119]. Hence, the gel-like behaviour exhibited by FUS-PLD droplets is not expected to occur in A-LCD-hnRNPA1 condensates with only one LARKS, since an inter-molecular network of β-sheets would not be able to fully percolate.

To further understand the gradual transformation of condensates during ageing from a molecular perspective, we now estimate the change to the free energy of the inter-molecular interaction network within the condensates due to ageing. To do so, we implement an energy-scaled molecular interaction analysis recently proposed in Refs. [87, 126]. Such analysis estimates the probability of contacts among all possible amino acid pairs in the condensate by considering not only a standard cut-off distance, but also the identity of the interacting amino acids via the mean excluded volume of the pair and the minimum potential energy of their interaction (both parameters taken from the coarse-grained force field). From the difference in the energy-scaled contact probability for pre-aged (*P_l_*) versus aged condensates (*P_g_*), we can estimate a free energy difference 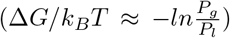 that directly relates to the transformation of the liquid-network connectivity as a result of ageing (further details on these calculations are provided in the Supplementary Methods)—which we term the ‘connectivity free energy difference’.

The connectivity free energy difference reveals that when condensate ageing is driven by the accumulation of inter-protein β-sheets, it is accompanied by a huge imbalance of inter-molecular forces. For instance, within A-LCD-hnRNPA1 and FUS-PLD condensates, LARKS–LARKS interactions give a large negative value of the connectivity free energy difference (approximately 5–6 k_*B*_T per residue), which evidences the long-lived nature of such connections as a result of ageing (Fig. 3a and Fig. 3b). In striking contrast, the connectivity free energy differences for the rest of amino acid pairs is negligible or even small and positive (Figs. 3a and Fig. 3b), consistent with weak and transient pre-ageing connections (i.e.,~0.5–1 k_*B*_T; Fig. 1a). In particular, the engagement of LARKS within structured β-sheet stacks significantly precludes their interaction with unstructured regions of neighbouring protein replicas, and results in moderate positive free energy differences. Such a severe imbalance in inter-molecular forces due to ageing—strong enough to drive the progressive dynamical arrest of proteins within droplets—contributes to rationalizing the physicochemical and molecular factors behind the intricate process of condensate ageing. Indeed, imbalance of inter-molecular forces has been shown to drive FUS single-component condensates to display multiphase architectures upon ageing [36, 127] or upon phosphorylation [86]. Furthermore, this imbalance is consistent with the formation of amorphous condensates observed in LARKS-containing proteins [58, 61, 63] such as hnRNPA1 [21], FUS [25], TDP-43 [128], or NUP-98 [33, 41].

### E. Ageing of condensates does not necessarily lead to loss of sphericity

We also explore how condensate ageing influences the shape of phase-separated droplets over time. First, we equilibrate a FUS-PLD condensate in non-ageing conditions (using the reference model [107, 108]) at T/T_*c,FUS*_=0.81 within a cubic box in the canonical ensemble (*NVT*: constant number of particles, volume and temperature). At this temperature, the phase-separated droplet displays liquid-like behaviour (Fig. 2a), and the condensed phase equilibrates into a roughly spherical droplet that minimizes the interfacial free energy of the system [129]. We quantify the sphericity of the droplet by computing the Solvent Available Surface Area (SASA) of the condensate (A) and its volume (*V*) for independent consecutive configurations. Through the following relation [130]: 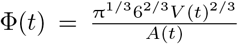, we can investigate the behaviour of the condensate sphericity (Φ) versus time. Sphericity values closer to 1, indicate that the condensate shapes approach a perfect sphere (further details on the order parameter to detect the biggest cluster and the SASA calculation are provided in the Supplementary Methods). In Figure 3d, we depict the average condensate sphericity for a liquid-like pre-aged condensate of 300 FUS-PLD protein replicas (red dashed line). When we activate our dynamical algorithm to trigger ageing, we observe that the droplet remains roughly spherical even at timescales where more than 75% of LARKS have transitioned into forming inter-protein β-sheets, and the condensate has become gradually kinetically-arrested (black curve; Fig. 3d). Our results, hence, predict that ageing of an already well-formed spherical condensate (i.e. that is not growing further due to droplet coalesence) would lead to negligible shape deformations. This is in agreement with previous simulations revealing that the origin of the widely recognised asphericity of aged condensates [25, 29] is non-ergodic droplet coalescence [37]. Fusion of small protein clusters to aged condensates is expected to drive the deformation of spherical condensates during ageing [37]. Moreover, impaired exchange of molecules between condensates and their surroundings, as observed in different multivalent proteins [131, 132], can lead to the emergence of irregular morphologies. However, reshaping of spherical liquid droplets during ageing, seems to play a negligible role in condensate amorphization, as shown in Fig. 3d.

### F. RNA decelerates the rate of accumulation of inter-protein β-sheets

The stability and viscoelastic properties of RNA-binding protein condensates is expected to be sensitively affected by the presence of RNA in an intricate manner that likely depends on RNA concentration, structure, sequence, and chain length [20, 21, 87, 113, 133, 134]. For instance, while short single-stranded disordered RNA strands (~50 nucleotides) can severely reduce droplet viscosity (e.g. of condensates made of LAF-1, an RNA-binding protein found in P granules [54]), long RNAs can increase viscosity in a concentration-dependent manner [55, 56]. Such complex impact of RNA has been reported for numerous RNA-binding protein condensates—e.g., of FUS [135–137], hnRNPA1 [21, 32, 87, 138], TDP-43 [139–141], TAF-15 [113, 142] or EWSR1 [20, 113, 142]—whose ageing has been associated with neurodegenerative diseases [26, 143]. Motivated by these observations, in this section we investigate the effect of single-stranded RNA on the viscoelasticity of FUS and A-LCD-hnRNPA1 condensates during the process of ageing.

We start by performing simulations of condensates that can age progressively over time due to the accumulation of inter-protein β-sheets, as we have done in the preceding sections. The key difference now is that we add varying concentrations of poly-Uridine (polyU). We choose polyU as it has been employed widely as a model for disordered single-stranded RNA in *in vitro* experiments examining RNA-binding protein condensates [54, 113, 116]. Regarding FUS, we only focus on the full protein (i.e., 526 residues) since the shorter FUS-PLD is devoid of RNA-recognition motifs (RRMs) and Arginine-Glycine rich-regions (RGGs), and does not present significant associative interactions with RNA at physiological conditions [58, 87].

RNA is well known to induce a concentrationdependent reentrant behaviour [114–116] for a wide range of RNA-binding protein pre-aged condensates— including, FUS [113–116], Whi3 [56], G3BP1 [144], or LAF-1 [54]. That is, low RNA concentrations moderately increase the stability of the condensates, while high RNA concentrations destabilize phase-separated droplets and solubilize them [113]. The specific RNA concentration thresholds at which the different behaviours emerge depend not only on the protein identity but also on the length of the RNA strand [87]. Here, we use a fixed RNA chain length of 125-nucleotides (nt), which is the minimum value for which moderate RNA concentrations can still enhance phase separation of FUS and A-LCD-hnRNPA1 condensates, as shown in our previous simulations [87]. For A-LCD-hnRNPA1, we showed that the optimal concentration of 125-nt polyU that maximizes the stability of the condensates (within the HPS-Cation-π model parameters) is 0.12 mg of polyU per mg of protein, and that mass ratios above 0.2 trigger condensate dissolution [87]. Indeed, beyond an RNA concentration of ~0.3 mg of polyU per mg of protein, A-LCD-hnRNPA1 phase-separated droplets fully dissolve at the studied temperature (and system size). For full FUS, in Figures 4a and 4b, we show the reentrant behaviour of pre-aged FUS condensates, as recapitulated by the HPS-Cation-π model [107, 108] in combination with the HPS-compatible model for RNA [111]. The phase diagrams of pre-aged FUS condensates show that, within such model parameters, concentrations of ~0.1 mg of polyU per mg of protein enhance FUS droplet stability, while higher concentrations of 0.2 mg of polyU per mg of protein reduce it.

**Figure 4:**
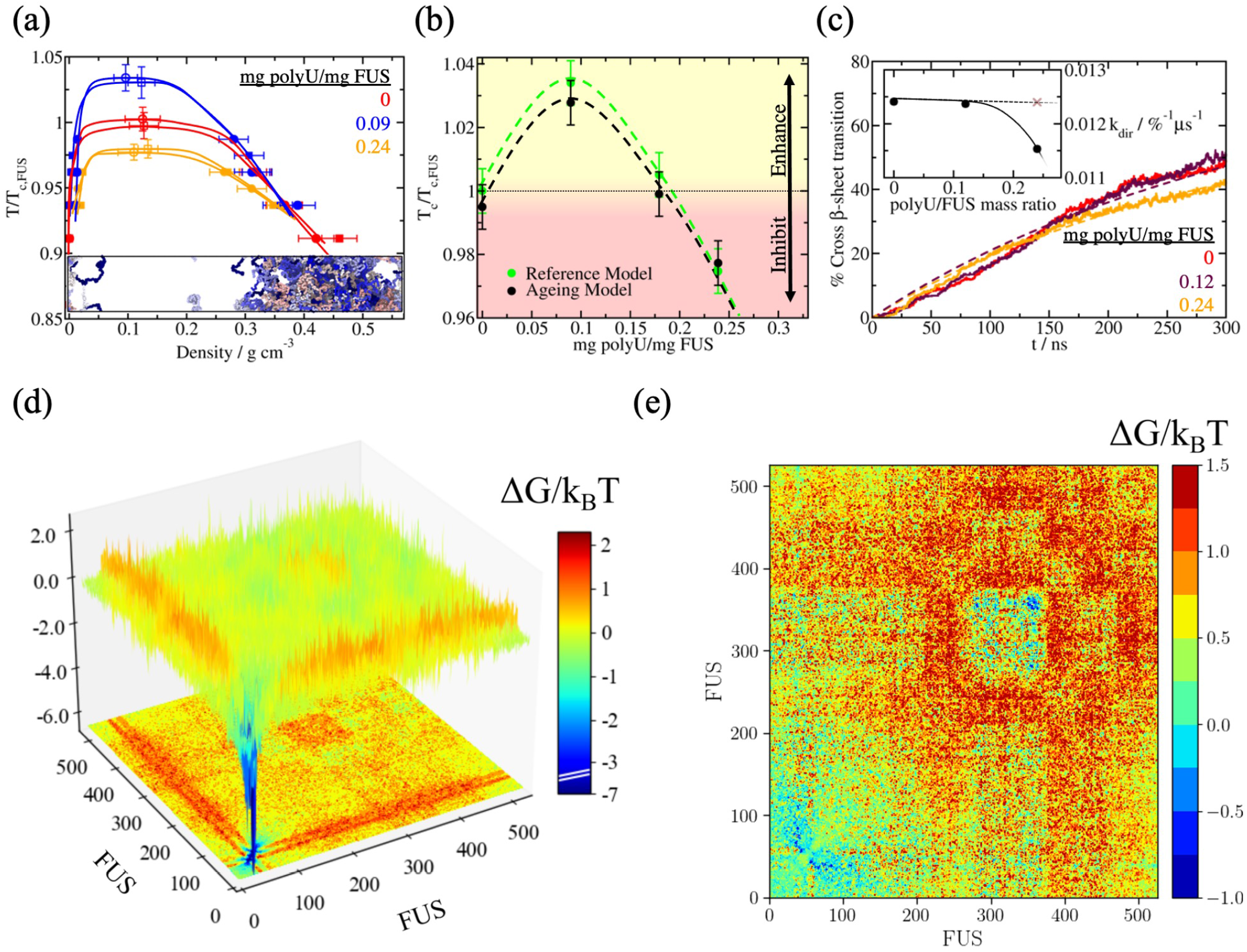
FUS disorder-to-order transitions are hindered by RNA which collectively contributes to blocking protein high-density fluctuations. (a) Temperature-density phase diagram for pure FUS (red symbols) and two polyU/FUS mixtures with different RNA concentrations as indicated in the legend. Filled circles represent the coexistence densities before structural transitions take place (reference model), and square symbols depict densities after ageing occurs (ageing model, i.e., >70% of LARKS within the condensates are engaged in inter-protein β-sheet motifs). Empty symbols indicate the estimated critical points obtained through the law of rectilinear diameters and critical exponents [110]. Please note that temperatures are normalized by the critical T of pure FUS (T*_c,FUS_*). Statistical errors are obtained by bootstrapping results from n = 3 independent simulations. (b) Critical temperature of FUS/polyU mixtures as a function of the polyU/FUS mass ratio evaluated for the reference model (green circles) and for the dynamical ageing model (black circles). Symbols above the horizontal dotted line imply LLPS enhancement while those below indicate phase-separation hindrance. The statistical uncertainty shown in Panel (a) also applies for (b). (c) Time-evolution of inter-protein β-sheet transitions (in percentage) within bulk condensates at different polyU/FUS mass ratios and at T/T_*c,FUS*_=0.96. Dashed lines account for second-order reaction fits to our data employed to estimate the kinetic constant of inter-protein β-sheet formation at distinct RNA concentrations. Inset: Structural transition kinetic constants as a function of polyU/FUS mass ratio within bulk phase-separated droplets. The brown cross depicts the computed kinetic constant in presence of inert polymers of the same length and concentration than polyU strands (black symbols). (d) Landscape of the average protein contact free energy variation upon condensate ageing of FUS droplets in absence of RNA at T/T_*c,FUS*_=0.96. ΔG/k_*B*_T is computed from the residue contact probability ratio between aged droplets (dynamical model) and liquid-like droplets (reference model). (e) Free energy inter-molecular variation computed from the molecular contact probability of pure FUS aged condensates and polyU/FUS (at 0.24 polyU/FUS mass ratio) aged condensates at T=0.96T_*c,FUS*_ and after an ageing time interval of ~1*μ*s for both systems.

We now investigate the consequences of adding RNA to condensates that age over time, due to the accumulation of inter-protein β-sheets. Consistent with our findings from Fig. 2a, the phase diagrams of FUS condensates and polyU/FUS droplets barely change during ageing (Fig. 4a; square symbols). Only a minor densification of the condensates over time is found when a very high percentage of β-sheet transitions has occurred (i.e., 75% of the available disorder LARKS have transitioned into inter-protein β-sheets). However, such modest increment of density [43], compared to the more prominent variation found in pure A-LCD-hnRNPA1 and FUS-PLD aged droplets (Figs. 1d and 2a respectively) is due to the small region in which the three LARKS are located within the full-FUS sequence (50-residue region within the 526-residue whole FUS sequence). To further illustrate the slight variation upon ageing in the phase diagram of polyU/FUS mixtures, we plot in Fig. 4b the critical temperature of FUS as a function of polyU/FUS mass ratio for condensates that have not yet aged (green circles; reference model), as well as for droplets that have already aged over time (black circles; ageing model). Like their pre-aged counterparts, aged condensates exhibit RNA-concentration driven reentrant behaviour. However, we also note that aged gel-like condensates with a high concentration of inter-protein β-sheet motifs (as those shown in Fig. 3c Left panel for FUS-PLD) may remain aggregated until reaching slightly higher temperatures than the critical one, hence showing moderate thermal hysteresis [37, 45, 145].

For both, condensates made of either A-LCD-hnRNPA1 or FUS, we find that their RNA-driven reentrant phase behaviour dictates the manner in which RNA impacts their ageing processes. Adding low concentrations of polyU (i.e., 0.09 mg of polyU per mg of protein for A-LCD-hnRNPA1 and 0.12 mg of polyU per mg of protein for FUS) has a negligible effect on the rate of accumulation of inter-protein β-sheets over time (Figs. 4c and 5a). However, as we go to higher RNA concentrations that approach the values needed to trigger condensate dissolution (e.g., 0.18 mg of polyU per mg of protein for A-LCD-hnRNPA1 and 0.24 mg of polyU per mg of protein for FUS), we observe a notable deceleration in the accumulation of inter-protein β-sheets for A-LCD-hnRNPA1 (Fig. 5a (Bottom panel); maroon curve) and a moderate reduction for FUS (Fig. 4c; orange curve). At high RNA concentrations, RNA–RNA electrostatic repulsion and steric hindrance begin to dominate, decreasing the condensate density, and as such, the probability of crucial high protein density fluctuations that enable the formation of inter-protein β-sheets [33, 35, 39]. Such effects are more modest for FUS ageing, because of its much larger size with respect to that of the LARKS-containing region, and because the main RNA-interacting domains in FUS (i.e., RRM and RGGs) are distantly located from the three LARKS motifs in the sequence. Nevertheless, condensates with higher RNA concentration present lower viscosities as a function of time (as shown in Fig. 5d for A-LCD-hnRNPA1/polyU condensates; maroon curve).

**Figure 5:**
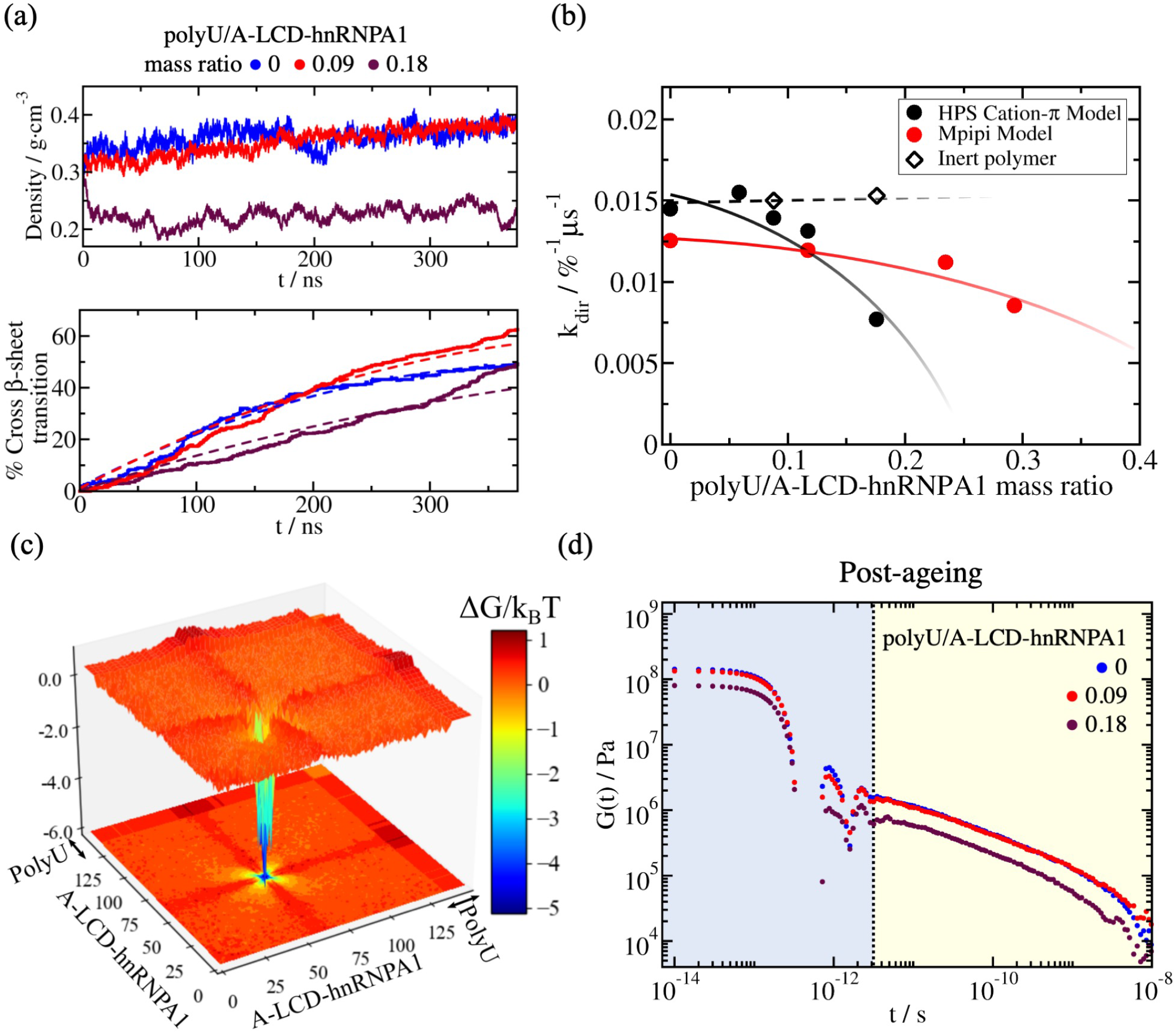
A-LCD-hnRNPA1 droplet ageing driven by disorder-to-order transitions is decelerated by inclusion of high polyU RNA concentration. (a) Time-evolution of droplet density (Top) and percentage of inter-protein β-sheet transitions within the condensates (Bottom) measured at different polyU/A-LCD-hnRNPA1 mass ratios and at T=0.97T_*c*_ (where T_*c*_ refers to the critical temperature of the pure protein condensate). Dashed lines in the bottom panel depict the second-order reaction fits employed to evaluate the kinetic constant (*k_dir_*) of inter-protein β-sheet formation (see details on the Supplementary Methods). (b) Estimated kinetic constants from a second-order reaction analysis to the number of inter-protein β-sheet transitions over time for different polyU/A-LCD-hnRNPA1 mass ratios at T=0.97T_*c*_ using two different residue-resolution models: HPS-Cation-π (black circles, [107, 108, 111]) and Mpipi (red circles, [112]). Black empty diamonds account for test control simulations in presence of inert polymers of the same length and concentration than polyU strands in the protein mixtures using the HPS-Cation-π model. Symbol sizes account for the estimated uncertainty while dotted and continuous lines are included as a guide for the eye. (c) Landscape of the protein and polyU (wide band) contacts free energy variation upon condensate ageing measured in polyU/A-LCD-hnRNPA1 phase-separated bulk droplets at T=0.97T*_c_* and 0.18 polyU/A-LCD-hnRNPA1 mass ratio. The same timescale for observing condensate ageing in pure A-LCD-hnRNPA1 droplets shown in Fig. 3b was explored here. (d) Shear stress relaxation modulus of polyU/A-LCD-hnRNPA1 aged condensates at T=0.97T_*c*_ for different polyU/protein mass ratios. The time interval to observe structural transitions before quenching them through our dynamical algorithm and compute *G*(*t*) was the same for all concentrations, ~1 *μ*s.

We now employ a simple chemical kinetic model to characterize in more detail the time-evolution of accumulation of inter-protein β-sheets within A-LCD-hnRNPA1 and FUS ageing condensates as a function of polyU concentration (at constant temperature of T=0.97T_*c*_ for A-LCD-hnRNPA1 and T=0.96T_*c,FUS*_ for FUS). For this, we use a second-order reaction analysis including two forward reactions:

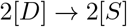

and

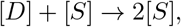

and one backward reaction:

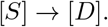

Here, [*D*] represents the percentage of fully disordered LARKS and [S] the percentage of LARKS forming inter-protein β-sheets. Through this analysis, we can estimate the kinetic constant *k_dir_* of inter-protein β-sheet formation (further details provided in the Supplementary Methods). For both A-LCD-hnRNPA1 (Fig. 5b) and FUS (Fig. 4c; inset), *k_dir_* significantly decreases as the polyU concentration is raised. For the reason described above, we again note that the effect is more modest for FUS than A-LCD-hnRNPA1. As a negative control, we compare the effects of inert polymers versus those of RNA. The inert polymers are composed of beads which exclusively exhibit excluded volume interactions (i.e., hard-spheres) and have the same size of our Uridine nucleotide beads. We set the concentrations of these inert polymers to match the monomeric concentrations of our previous RNA/protein mixtures. As depicted in Fig. 4c (brown crosses) and Fig. 5b (empty diamonds) for FUS and A-LCD-hnRNPA1 respectively, the accumulation of structural transitions is faster (i.e., *k_dir_* is significantly higher) when polyU strands are substituted by inert polymers, and is of the same order as in the pure protein condensates. Moreover, the addition of shorter disordered proteins (also at similar monomeric concentrations as RNA) that could contribute to destabilizing biomolecular condensates and reducing their density due to the lower critical temperature of the shorter proteins (i.e., as FUS-PLDs mixed in full-FUS condensates; Fig. 2a), moderately increases the rate of β-sheet transitions over time as compared to pure full-FUS systems (Supplementary Figure 3). These results evidence the specific ability of RNA to decelerate ageing driven by accumulation of inter-protein β-sheets inside condensates. Our simulations reveal that disorder-to-order transitions within condensates (triggered by the spontaneous formation of high density protein clusters) are inhibited by RNA via the combination of two factors: (1) binding of RNA to various protein regions (i.e., RRMs or RGGs), which reduces the likelihood of inter-protein interactions, and (2) the RNA–RNA long-range electrostatic repulsion, which critically lowers the condensate density. Since inert polymers or small disordered proteins (like FUS-PLD) cannot fulfill either of these two roles that RNA accomplishes at high concentration, they cannot decelerate condensate ageing at similar monomeric concentrations (Figs. 4c (inset) and 5b). Additionally, to demonstrate that our observations for polyU/protein mixtures are not model-dependent, we repeat the calculations for A-LCD-hnRNPA1 using the Mpipi force field residue resolution model [112], which recapitulates quantitatively the temperature–density experimental phase diagram of such protein [112]. Using the Mpipi (Fig. 5b; red circles), we also find that the kinetic constant k_dir_ decreases with the concentration of RNA; hence, delaying condensate gelation over time (Fig. 5d).

Comparing how the energy-scaled molecular interactions (see Supplementary Methods) change upon ageing in the absence and presence of RNA, provides microscopic insight on such modulation. While, in the absence of RNA, the A-LCD-hnRNAP1 ageing connectivity free energy difference per residue is large for LARKS–LARKS interactions (i.e., ~·6k_*B*_T for A-LCD-hnRNPA1), at high RNA concentrations its value decreases to ~4.4k_*B*_T for the same interactions (Fig. 5c). Such a significant reduction explains the shorter relaxation times and lower viscosity of A-LCD-hnRNPA1 condensates with high concentrations of polyU (Fig. 5d). Indeed, previous simulations have shown that variations of the order of 1k_*B*_T in protein binding can transform protein self-diffusion by several orders of magnitude [37]. Our findings therefore clarify from a mechanistic and molecular perspective previous experimental results showing that: (1) hnRNPA1 fibrillization is enhanced in protein-rich droplets formed via liquid-liquid phase separation [21, 32, 138]; and (2) liquid-to-solid aberrant phase transitions, here driven by inter-protein β-sheet transitions, might be prevented by keeping RNA-binding proteins soluble at high RNA concentrations [113].

For full-FUS pure condensates, the average binding free energy gain associated with LARKS–LARKS β-sheets is of approximately 5k_*B*_T per residue. Moreover, due to the local binding strengthening of LARKS– LARKS interactions within the PLD, other regions within the FUS sequence (such as RGGs, RMMs or the zinc finger), moderately increase their binding probability by ~0.5k_*B*_T (Fig. 4d; light green regions). Hence, FUS phase-separated droplets collectively boost their enthalpic gain during ageing via the progressive accumulation of inter-protein β-sheet structures [26, 33, 35, 36]. However, if we analyse the variance of the aged system when exposed to a high concentration of RNA, such protein–protein interaction gain vanishes, and is substituted by more favourable protein–RNA interactions, especially between RNA–RRM and RNA–RGG domains. Thus, during FUS ageing, the net variation in protein binding probability upon RNA inclusion is positive, and ΔG/k_*B*_T increases by almost ~1.5k_*B*_T on average for most of the sequence regions (implying lower protein binding probability; Fig. 4e). Such free energy increase collectively hinders the high-density protein local fluctuations within FUS/RNA condensates that underlie the progressive emergence of inter-protein β-sheet transitions [35, 38, 39, 104]. This occurs despite RNA binding not targeting directly the FUS-PLD region (1-163 residues; where the binding strength decreases by ~0.15k_*B*_T), which is the ageing epicentre of FUS.

## III. DISCUSSION

In this work, we develop an innovative multiscale computational approach that integrates all-atom simulations and sequence-dependent coarse-grained models to microscopically elucidate the progressive ageing of protein condensates due to inter-protein structural changes. First, we show how the accumulation of inter-protein LARKS β-sheets, can critically enhance inter-molecular protein binding to drive liquid-to-gel transitions in biomolecular condensates. We find that the reorganization of A-LCD-hnRNPA1 disordered LARKS into structured four-peptide β-sheet fibrils can transform weak and transient protein interactions into almost irreversible long-lived contacts. Such local binding strengthening, even in the absence of chemical modifications or external stimuli, can dramatically increase condensate viscosity and moderately raise droplet density, as we report here for A-LCD-hnRNPA1 and FUS condensates. Our findings from this study may also explain how aberrant phase transitions in other LARKS-containing proteins such as TDP-43, Aβ-NKGAII or NUP-98 among many others [33, 35, 37, 39] can be regulated by different factors such as temperature, RNA or protein concentration. Moreover, we observe that ageing driven by accumulation of inter-peptide β-sheet transitions critically alters the molecular network of connections sustaining the condensate. Our results suggest that the widely recognised asphericity as a consequence of condensate ageing [25, 29, 146], emerges indirectly during ageing, from non-ergodic droplet coalescence [37] rather than directly from ageing condensate reshaping (Fig. 3d).

Remarkably, we also find that recruitment of high concentrations of RNA by condensates significantly slows down the rate of accumulation (i.e., kinetic constant) of inter-protein β-sheets over time. Such reduction in the kinetic constant, which is not observed at low or moderate RNA concentrations, has a critical impact in ensuring droplet viscoelastic properties remain consistent with those of liquids rather than of gels during ageing. Hence, our results suggest that high RNA concentrations may contribute to reducing the onset of pathological liquid-to-solid transitions during ageing [20, 26]. The decrease in: (1) protein binding probability induced by the emergence of favourable protein-RNA interactions, and (2) condensate density due to RNA-RNA electrostatic repulsion, collectively contributes to frustrating the high-density protein fluctuations that would otherwise promote inter-peptide β-sheet formation. However, these conditions may only be satisfied beyond the optimal RNA concentration enhancing LLPS (i.e., RNA reentrant point [114–116]). Taken together, our multiscale simulations shed light on the physicochemical and molecular factors behind the intricate process of pathological ageing in LARKS-containing proteins—such as FUS [25] or hnRNPA1 [21] among others [33, 41]—and suggest a potential framework to decelerate aberrant phase transitions due to protein structural changes.

## Supporting information

Description of Supplementary Movies

## ACKNOWLEDGMENTS

This project has received funding from the Oppenheimer Research Fellowship of the University of Cambridge. A. T. is funded by Universidad Politécnica de Madrid (ESTANCIAS-PIF 20-TYOSR8-13-4N0WPQ and PhD fellowship ‘programa propio UPM’) and the Oppenheimer Fellowship, I. S.-B. acknowledges funding from the Oppenheimer Fellowship, Derek Brewer scholarship of Emmanuel College and EPSRC Doctoral Training Programme studentship, number EP/T517847/1. A. G. is funded by an EPSRC studentship (EP/N509620/1) and a Winton scholarship. J. R. acknowledges funding from the Spanish Ministry of Economy and Competitivity (PID2019-105898GA-C22) and the Madrid Government (Comunidad de Madrid-Spain) under the Multiannual Agreement with Universidad Politécnica de Madrid in the line Excellence Programme for University Professors, in the context of the V PRICIT (Regional Programme of Research and Technological Innovation). J. R. E. also acknowledges funding from the Roger Ekins Research Fellowship of Emmanuel College. M. E. acknowledges funding from the Oppenheimer Fellowship. This work has been performed using resources provided by the Cambridge Tier-2 system operated by the University of Cambridge Research Computing Service (http://www.hpc.cam.ac.uk) funded by EPSRC Tier-2 capital grant EP/P020259/1. The authors gratefully acknowledge the Universidad Politécnica de Madrid (www.upm.es) for also providing computing resources on Magerit Supercomputer. We also acknowledge critical reading of the paper by V. Roser and helpful discussions with J. Cortijo.

## DATA AVAILABILITY

The data that supports the findings of this study are available within the article and its Supplementary Methods. The LAMMPS and GROMACS files of the residue-resolution models and all-atom simulations respectively, as well as the dynamic algorithm software are available in the GitHub database under the accession code: [https://doi.org/10.5281/zenodo.6979617]. The following PDB files can be obtained through these codes: PDB Code 6BXX [http://doi.org/10.2210/pdb6bxx/pdb], PDB Code 2LCW [http://doi.org/10.2210/pdb2lcw/pdb], and PDB Code 6G99 [http://doi.org/10.2210/pdb6g99/pdb].

## AUTHOR CONTRIBUTIONS

J.R. and J.R.E designed research; A.R.T., I.S.-B. and M.E.-E. performed research; A.R.T., I.S.-B. and M.E.-E. analyzed data; A.R.T., I.S.-B., A.G., R.C.-G., J.R. and J.R.E. developed new methods; J.R. and J.R.E. supervised research; and R.C.-G. and J.R.E. wrote the original draft. All authors edited the paper.

## References

[1] Brangwynne, C. P. et al. Germline p granules are liquid droplets that localize by controlled dissolution/condensation. Science 324, 1729–1732 (2009).

[2] Alberti, S., Gladfelter, A. & Mittag, T. Considerations and challenges in studying liquid-liquid phase separation and biomolecular condensates. Cell 176, 419–434 (2019).

[3] Li, P. et al. Phase transitions in the assembly of multivalent signalling proteins. Nature 483, 336–340 (2012).

[4] Hyman, A. A., Weber, C. A. & Jülicher, F. Liquid-liquid phase separation in biology. Annual review of cell and developmental biology 30, 39–58 (2014).

[5] Labbé, K., Murley, A. & Nunnari, J. Determinants and functions of mitochondrial behavior. Annual review of cell and developmental biology 30, 357–391 (2014).

[6] Solovei, I., Thanisch, K. & Feodorova, Y. How to rule the nucleus: divide et impera. Current opinion in cell biology 40, 47–59 (2016).

[7] Rothman, J. E. The golgi apparatus: two organelles in tandem. Science 213, 1212–1219 (1981).

[8] Banani, S. F. et al. Compositional control of phase-separated cellular bodies. Cell 166, 651–663 (2016).

[9] Shin, Y. & Brangwynne, C. P. Liquid phase condensation in cell physiology and disease. Science 357, eaaf4382 (2017).

[10] Nott, T. J. et al. Phase transition of a disordered nuage protein generates environmentally responsive membraneless organelles. Molecular cell 57, 936–947 (2015).

[11] Narlikar, G. J. Phase-separation in chromatin organization. Journal of Biosciences 45, 5 (2020).

[12] Brangwynne, C. P., Tompa, P. & Pappu, R. V. Polymer physics of intracellular phase transitions. Nature Physics 11, 899–904 (2015).

[13] Protter, D. S. et al. Intrinsically disordered regions can contribute promiscuous interactions to rnp granule assembly. Cell reports 22, 1401–1412 (2018).

[14] Sanders, D. W. et al. Competing protein-rna interaction networks control multiphase intracellular organization. Cell 181, 306–324 (2020).

[15] Yoo, H., Triandafillou, C. & Drummond, D. A. Cellular sensing by phase separation: Using the process, not just the products. Journal of Biological Chemistry 294, 7151–7159 (2019).

[16] Riback, J. A. et al. Innovative scattering analysis shows that hydrophobic disordered proteins are expanded in water. Science 358, 238–241 (2017).

[17] Krainer, G. et al. Reentrant liquid condensate phase of proteins is stabilized by hydrophobic and non-ionic interactions. Nature Communications 12, 1–14 (2021).

[18] Xiao, Q., McAtee, C. K. & Su, X. Phase separation in immune signalling. Nature Reviews Immunology 22, 1–12 (2021).

[19] Franzmann, T. M. et al. Phase separation of a yeast prion protein promotes cellular fitness. Science 359, eaao5654 (2018).

[20] Guo, L. & Shorter, J. It’s raining liquids: Rna tunes viscoelasticity and dynamics of membraneless organelles. Molecular cell 60, 189–192 (2015).

[21] Molliex, A. et al. Phase separation by low complexity domains promotes stress granule assembly and drives pathological fibrillization. Cell 163, 123–133 (2015).

[22] Patel, A. et al. A liquid-to-solid phase transition of the als protein fus accelerated by disease mutation. Cell 162, 1066–1077 (2015).

[23] Rabouille, C. & Alberti, S. Cell adaptation upon stress: the emerging role of membrane-less compartments. Current opinion in cell biology 47, 34–42 (2017).

[24] Monahan, Z. et al. Phosphorylation of the fus low-complexity domain disrupts phase separation, aggregation, and toxicity. The EMBO journal 36, 2951–2967 (2017).

[25] Jawerth, L. et al. Protein condensates as aging maxwell fluids. Science 370, 1317–1323 (2020).

[26] Portz, B., Lee, B. L. & Shorter, J. Fus and tdp-43 phases in health and disease. Trends in Biochemical Sciences 46, 550–563 (2021).

[27] Vance, C. et al. Mutations in fus, an rna processing protein, cause familial amyotrophic lateral sclerosis type 6. Science 323, 1208–1211 (2009).

[28] Kamelgarn, M. et al. Als mutations of fus suppress protein translation and disrupt the regulation of nonsense-mediated decay. Proceedings of the National Academy of Sciences 115, E11904–E11913 (2018).

[29] Ray, S. et al. *α*-synuclein aggregation nucleates through liquid–liquid phase separation. Nature chemistry 12, 705–716 (2020).

[30] Wegmann, S. et al. Tau protein liquid–liquid phase separation can initiate tau aggregation. The EMBO Journal 37, e98049 (2018).

[31] Alberti, S. & Dormann, D. Liquid–liquid phase separation in disease. Annual Review of Genetics 53, 171–194 (2019).

[32] Gui, X. et al. Structural basis for reversible amyloids of hnrnpa1 elucidates their role in stress granule assembly. Nature communications 10, 1–12 (2019).

[33] Hughes, M. P. et al. Atomic structures of low-complexity protein segments reveal kinked sheets that assemble networks. Science 359, 698 (2018).

[34] Sun, Y. et al. The nuclear localization sequence mediates hnrnpa1 amyloid fibril formation revealed by cryoem structure. Nature communications 11, 1–8 (2020).

[35] Luo, F. et al. Atomic structures of fus lc domain segments reveal bases for reversible amyloid fibril formation. Nature Structural & Molecular Biology 25, 341–346 (2018).

[36] Garaizar, A. et al. Aging can transform single-component protein condensates into multiphase architectures. Proceedings of the National Academy of Sciences 119, e2119800119 (2022).

[37] Garaizar, A., Espinosa, J. R., Joseph, J. A. & Collepardo-Guevara, R. Kinetic interplay between droplet maturation and coalescence modulates shape of aged protein condensates. Scientific reports 12, 1–13 (2022).

[38] Šarić, A., Chebaro, Y. C., Knowles, T. P. & Frenkel, D. Crucial role of nonspecific interactions in amyloid nucleation. Proceedings of the National Academy of Sciences 111, 17869–17874 (2014).

[39] Guenther, E. L. et al. Atomic structures of tdp-43 lcd segments and insights into reversible or pathogenic aggregation. Nature Structural & Molecular Biology 25, 463–471 (2018).

[40] Fonda, B. D., Jami, K. M., Boulos, N. R. & Murray, D. T. Identification of the rigid core for aged liquid droplets of an rna-binding protein low complexity domain. Journal of the American Chemical Society 143, 6657–6668 (2021).

[41] Milles, S. et al. Facilitated aggregation of fg nucleoporins under molecular crowding conditions. EMBO reports 14, 178–183 (2013).

[42] Mendoza-Espinosa, P., García-González, V., Moreno, A., Castillo, R. & Mas-Oliva, J. Disorder-to-order conformational transitions in protein structure and its relationship to disease. Molecular and cellular biochemistry 330, 105–120 (2009).

[43] Chatterjee, S. et al. Reversible kinetic trapping of fus biomolecular condensates. Advanced Science 9, 2104247 (2022).

[44] Kato, M. et al. Cell-free formation of rna granules: low complexity sequence domains form dynamic fibers within hydrogels. Cell 149, 753–767 (2012).

[45] Murakami, T. et al. Als/ftd mutation-induced phase transition of fus liquid droplets and reversible hydrogels into irreversible hydrogels impairs rnp granule function. Neuron 88, 678–690 (2015).

[46] Hennig, S. et al. Prion-like domains in rna binding proteins are essential for building subnuclear paraspeckles. Journal of Cell Biology 210, 529–539 (2015).

[47] Liu, C. et al. Out-of-register-sheets suggest a pathway to toxic amyloid aggregates. Proceedings of the National Academy of Sciences 109, 20913–20918 (2012).

[48] Ambadipudi, S., Biernat, J., Riedel, D., Mandelkow, E. & Zweckstetter, M. Liquid–liquid phase separation of the microtubule-binding repeats of the alzheimer-related protein tau. Nature Communications 8, 275 (2017).

[49] Chuang, E., Hori, A. M., Hesketh, C. D. & Shorter, J. Amyloid assembly and disassembly. Journal of Cell Science 131, jcs189928 (2018).

[50] Banani, S. F., Lee, H. O., Hyman, A. A. & Rosen, M. K. Biomolecular condensates: organizers of cellular biochemistry. Nature reviews Molecular cell biology 18, 285–298 (2017).

[51] Lin, Y., Protter, D. S., Rosen, M. K. & Parker, R. Formation and Maturation of Phase-Separated Liquid Droplets by RNA-Binding Proteins. Molecular Cell 60, 208–219 (2015).

[52] Jang, S. et al. Phosphofructokinase relocalizes into subcellular compartments with liquid-like properties in vivo. Biophysical Journal 120, 1170–1186 (2020).

[53] Nair, S. J. et al. Phase separation of ligand-activated enhancers licenses cooperative chromosomal enhancer assembly. Nature Structural & Molecular Biology 26, 193–203 (2019).

[54] Elbaum-Garfinkle, S. et al. The disordered p granule protein laf-1 drives phase separation into droplets with tunable viscosity and dynamics. Proceedings of the National Academy of Sciences 112, 7189–7194 (2015).

[55] Wei, M.-T. et al. Phase behaviour of disordered proteins underlying low density and high permeability of liquid organelles. Nature Chemistry 9, 1118 (2017).

[56] Zhang, H. et al. Rna controls polyq protein phase transitions. Molecular cell 60, 220–230 (2015).

[57] Michieletto, D. & Marenda, M. Rheology and viscoelasticity of proteins and nucleic acids condensates. JACS Au 2, 1506–1521 (2022).

[58] Wang, A. et al. A single n-terminal phosphomimic disrupts tdp-43 polymerization, phase separation, and rna splicing. The EMBO journal 37, e97452 (2018).

[59] March, Z. M., King, O. D. & Shorter, J. Prion-like domains as epigenetic regulators, scaffolds for subcellular organization, and drivers of neurodegenerative disease. Brain research 1647, 9–18 (2016).

[60] Gotor, N. L. et al. Rna-binding and prion domains: the yin and yang of phase separation. Nucleic acids research 48, 9491–9504 (2020).

[61] Murray, D. T. et al. Structure of fus protein fibrils and its relevance to self-assembly and phase separation of low-complexity domains. Cell 171, 615–627 (2017).

[62] Alberti, S. & Hyman, A. A. Biomolecular condensates at the nexus of cellular stress, protein aggregation disease and ageing. Nature Reviews Molecular Cell Biology 22, 196–213 (2021).

[63] Shen, Y. et al. Biomolecular condensates undergo a generic shear-mediated liquid-to-solid transition. Nature nanotechnology 15, 841–847 (2020).

[64] Wen, J. et al. Conformational expansion of tau in condensates promotes irreversible aggregation. Journal of the American Chemical Society 143, 13056–13064 (2021).

[65] Mathieu, C., Pappu, R. V. & Taylor, J. P. Beyond aggregation: Pathological phase transitions in neurodegenerative disease. Science 370, 56–60 (2020).

[66] Dignon, G. L., Best, R. B. & Mittal, J. Biomolecular phase separation: From molecular driving forces to macroscopic properties. Annual review of physical chemistry 71, 53–75 (2020).

[67] Schuster, B. S. et al. Identifying sequence perturbations to an intrinsically disordered protein that determine its phase-separation behavior. Proceedings of the National Academy of Sciences 117, 11421–11431 (2020).

[68] Welsh, T. J. et al. Surface electrostatics govern the emulsion stability of biomolecular condensates. Nano letters 2, 612–621 (2022).

[69] Paloni, M., Bailly, R., Ciandrini, L. & Barducci, A. Unraveling molecular interactions in liquid–liquid phase separation of disordered proteins by atomistic simulations. The Journal of Physical Chemistry B 124, 9009–9016 (2020).

[70] Zheng, W. et al. Molecular details of protein condensates probed by microsecond long atomistic simulations. The Journal of Physical Chemistry B 124, 11671–11679 (2020).

[71] Dignon, G. L., Zheng, W., Best, R. B., Kim, Y. C. & Mittal, J. Relation between single-molecule properties and phase behavior of intrinsically disordered proteins. Proceedings of the National Academy of Sciences of the United States of America 115, 9929–9934 (2018).

[72] Dignon, G. L., Zheng, W., Kim, Y. C. & Mittal, J. Temperature-controlled liquid–liquid phase separation of disordered proteins. ACS central science 5, 821–830 (2019).

[73] Garaizar, A. & Espinosa, J. R. Salt dependent phase behavior of intrinsically disordered proteins from a coarsegrained model with explicit water and ions. The Journal of Chemical Physics 155, 125103 (2021).

[74] Sanchez-Burgos, I., Joseph, J. A., Collepardo-Guevara, R. & Espinosa, J. R. Size conservation emerges spontaneously in biomolecular condensates formed by scaffolds and surfactant clients. Scientific Reports 11, 1–10 (2021).

[75] Benayad, Z., von Bülow, S., Stelzl, L. S. & Hummer, G. Simulation of fus protein condensates with an adapted coarse-grained model. Journal of Chemical Theory and Computation 17, 525–537 (2020).

[76] Harmon, T. S., Holehouse, A. S., Rosen, M. K. & Pappu, R. V. Intrinsically disordered linkers determine the interplay between phase separation and gelation in multivalent proteins. elife 6, e30294 (2017).

[77] Garaizar, A., Sanchez-Burgos, I., Collepardo-Guevara, R. & Espinosa, J. R. Expansion of intrinsically disordered proteins increases the range of stability of liquid– liquid phase separation. Molecules 25, 4705 (2020).

[78] Sanchez-Burgos, I., Espinosa, J. R., Joseph, J. A. & Collepardo-Guevara, R. Valency and binding affinity variations can regulate the multilayered organization of protein condensates with many components. Biomolecules 11, 278 (2021).

[79] Statt, A., Casademunt, H., Brangwynne, C. P. & Panagiotopoulos, A. Z. Model for disordered proteins with strongly sequence-dependent liquid phase behavior. The Journal of Chemical Physics 152, 075101 (2020).

[80] Das, S., Eisen, A., Lin, Y.-H. & Chan, H. S. A lattice model of charge-pattern-dependent polyampholyte phase separation. The Journal of Physical Chemistry B 122, 5418–5431 (2018).

[81] Choi, J. M., Dar, F. & Pappu, R. V. LASSI: A lattice model for simulating phase transitions of multivalent proteins. PLoS Computational Biology 15, e1007028 (2019).

[82] Jacobs, W. M. Self-assembly of biomolecular condensates with shared components. Physical review letters 126, 258101 (2021).

[83] Weber, C. A., Zwicker, D., Jülicher, F. & Lee, C. F. Physics of active emulsions. Reports on Progress in Physics 82, 064601 (2019).

[84] Wurtz, J. D. & Lee, C. F. Stress granule formation via atp depletion-triggered phase separation. New Journal of Physics 20, 045008 (2018).

[85] Weber, C. A., Lee, C. F. & Jülicher, F. Droplet ripening in concentration gradients. New Journal of Physics 19, 053021 (2017).

[86] Ranganathan, S. & Shakhnovich, E. Effect of rna on morphology and dynamics of membraneless organelles. The Journal of Physical Chemistry B 125, 5035–5044 (2021).

[87] Tejedor, A. R., Garaizar, A., Ramírez, J. & Espinosa, J. R. Rna modulation of transport properties and stability in phase separated condensates. Biophysical Journal 120, 5169–5186 (2021).

[88] Alshareedah, I., Moosa, M. M., Pham, M., Potoyan, D. A. & Banerjee, P. R. Programmable viscoelasticity in protein-rna condensates with disordered sticker-spacer polypeptides. Nature communications 12, 1–14 (2021).

[89] Alberti, S. & Hyman, A. A. Are aberrant phase transitions a driver of cellular aging? BioEssays 38, 959–968 (2016).

[90] Babinchak, W. M. & Surewicz, W. K. Liquid–liquid phase separation and its mechanistic role in pathological protein aggregation. Journal of Molecular Biology 432, 1910 – 1925 (2020).

[91] Nedelsky, N. B. & Taylor, J. P. Bridging biophysics and neurology: aberrant phase transitions in neurodegenerative disease. Nature Reviews Neurology 15, 272–286 (2019).

[92] Patel, A. et al. A liquid-to-solid phase transition of the als protein fus accelerated by disease mutation. Cell 162, 1066 – 1077 (2015).

[93] Spannl, S., Tereshchenko, M., Mastromarco, G. J., Ihn, S. J. & Lee, H. O. Biomolecular condensates in neurodegeneration and cancer. Traffic 20, 890–911 (2019).

[94] Pytowski, L., Lee, C. F., Foley, A. C., Vaux, D. J. & Jean, L. Liquid–liquid phase separation of type ii diabetes-associated iapp initiates hydrogelation and aggregation. Proceedings of the National Academy of Sciences 117, 12050 (2020).

[95] Bosco, D. A. et al. Mutant fus proteins that cause amyotrophic lateral sclerosis incorporate into stress granules. Human molecular genetics 19, 4160–4175 (2010).

[96] Li, Y. R., King, O. D., Shorter, J. & Gitler, A. D. Stress granules as crucibles of als pathogenesis. Journal of cell biology 201, 361–372 (2013).

[97] Espinosa, J. R. et al. Liquid network connectivity regulates the stability and composition of biomolecular condensates with many components. Proceedings of the National Academy of Sciences 117, 13238–13247 (2020).

[98] Lemkul, J. A. & Bevan, D. R. Assessing the stability of alzheimer’s amyloid protofibrils using molecular dynamics. The Journal of Physical Chemistry B 114, 1652–1660 (2010).

[99] Robustelli, P., Piana, S. & Shaw, D. E. Developing a molecular dynamics force field for both folded and disordered protein states. Proceedings of the National Academy of Sciences 115, E4758–E4766 (2018).

[100] Huang, J. et al. Charmm36m: an improved force field for folded and intrinsically disordered proteins. Nature methods 14, 71–73 (2017).

[101] Samantray, S., Yin, F., Kav, B. & Strodel, B. Different force fields give rise to different amyloid aggregation pathways in molecular dynamics simulations. Journal of chemical information and modeling 60, 6462–6475 (2020).

[102] Martin, E. W. et al. Interplay of folded domains and the disordered low-complexity domain in mediating hnrnpa1 phase separation. Nucleic acids research 49, 2931–2945 (2021).

[103] Mitrea, D. M. & Kriwacki, R. W. Phase separation in biology; functional organization of a higher order. Cell Communication and Signaling 14, 1–20 (2016).

[104] Michaels, T. C. et al. Chemical kinetics for bridging molecular mechanisms and macroscopic measurements of amyloid fibril formation. Annual review of physical chemistry 69, 273–298 (2018).

[105] Lu, Y., Lim, L. & Song, J. Rrm domain of als/ftd-causing fus characteristic of irreversible unfolding spontaneously self-assembles into amyloid fibrils. Scientific reports 7, 1–14 (2017).

[106] Ramírez-Alvarado, M., Merkel, J. S. & Regan, L. A systematic exploration of the influence of the protein stability on amyloid fibril formation in vitro. Proceedings of the National Academy of Sciences 97, 8979–8984 (2000).

[107] Das, S., Lin, Y.-H., Vernon, R. M., Forman-Kay, J. D. & Chan, H. S. Comparative roles of charge, π, and hydrophobic interactions in sequence-dependent phase separation of intrinsically disordered proteins. Proceedings of the National Academy of Sciences 117, 28795–28805 (2020).

[108] Dignon, G. L., Zheng, W., Kim, Y. C., Best, R. B. & Mittal, J. Sequence determinants of protein phase behavior from a coarse-grained model. PLoS computational biology 14, e1005941 (2018).

[109] Ladd, A. J. & Woodcock, L. V. Triple-point coexistence properties of the lennard-jones system. Chemical Physics Letters 51, 155–159 (1977).

[110] Rowlinson, J. S. & Widom, B. Molecular theory of capillarity (Courier Corporation, 2013).

[111] Regy, R. M., Dignon, G. L., Zheng, W., Kim, Y. C. & Mittal, J. Sequence dependent phase separation of protein-polynucleotide mixtures elucidated using molecular simulations. Nucleic Acids Research 48, 12593–12603 (2020).

[112] Joseph, J. A. et al. Physics-driven coarse-grained model for biomolecular phase separation with near-quantitative accuracy. Nature Computational Science 1, 732–743 (2021).

[113] Maharana, S. et al. Rna buffers the phase separation behavior of prion-like rna binding proteins. Science 360, 918–921 (2018).

[114] Burke, K. A., Janke, A. M., Rhine, C. L. & Fawzi, N. L. Residue-by-residue view of in vitro fus granules that bind the c-terminal domain of rna polymerase ii. Molecular cell 60, 231–241 (2015).

[115] Schwartz, J. C., Wang, X., Podell, E. R. & Cech, T. R. Rna seeds higher-order assembly of fus protein. Cell reports 5, 918–925 (2013).

[116] Banerjee, P. R., Milin, A. N., Moosa, M. M., Onuchic, P. L. & Deniz, A. A. Reentrant phase transition drives dynamic substructure formation in ribonucleoprotein droplets. Angewandte Chemie 129, 11512–11517 (2017).

[117] Ladd, A. & Woodcock, L. Triple-point coexistence properties of the lennard-jones system. Chemical Physics Letters 51, 155–159 (1977).

[118] Espinosa, J. R., Sanz, E., Valeriani, C. & Vega, C. On fluid-solid direct coexistence simulations: The pseudo-hard sphere model. The Journal of chemical physics 139, 144502 (2013).

[119] Rubinstein, M., Colby, R. H. et al. Polymer physics, vol. 23 (Oxford university press New York, 2003).

[120] Ramírez, J., Sukumaran, S. K., Vorselaars, B. & Likhtman, A. E. Efficient on the fly calculation of time correlation functions in computer simulations. The Journal of chemical physics 133, 154103 (2010).

[121] St George-Hyslop, P. et al. The physiological and pathological biophysics of phase separation and gelation of rna binding proteins in amyotrophic lateral sclerosis and fronto-temporal lobar degeneration. Brain Research 1693, 11–23 (2018). RNA Metabolism in Neurological Disease 2018.

[122] Ferrolino, M. C., Mitrea, D. M., Michael, J. R. & Kriwacki, R. W. Compositional adaptability in npm1-surf6 scaffolding networks enabled by dynamic switching of phase separation mechanisms. Nature Communications 9, 5064 (2018).

[123] Woodruff, J. B. et al. The centrosome is a selective condensate that nucleates microtubules by concentrating tubulin. Cell 169, 1066–1077.e10 (2017).

[124] Sukumaran, S. K., Grest, G. S., Kremer, K. & Everaers, R. Identifying the primitive path mesh in entangled polymer liquids. Journal of Polymer Science Part B: Polymer Physics 43, 917–933 (2005).

[125] Hagita, K. & Murashima, T. Effect of chain-penetration on ring shape for mixtures of rings and linear polymers. Polymer 218, 123493 (2021).

[126] Tesei, G., Schulze, T. K., Crehuet, R. & Lindorff-Larsen, K. Accurate model of liquid–liquid phase behavior of intrinsically disordered proteins from optimization of single-chain properties. Proceedings of the National Academy of Sciences 118, e2111696118 (2021).

[127] Erkamp, N. A. et al. Multiphase condensates from a kinetically arrested phase transition. bioRxiv (2022).

[128] Zhuo, X.-F. et al. Solid-state nmr reveals the structural transformation of the tdp-43 amyloidogenic region upon fibrillation. Journal of the American Chemical Society 142, 3412–3421 (2020).

[129] Montero de Hijes, P., Espinosa, J. R., Sanz, E. & Vega, C. Interfacial free energy of a liquid-solid interface: Its change with curvature. The Journal of Chemical Physics 151, 144501 (2019).

[130] Wadell, H. Volume, shape, and roundness of quartz particles. The Journal of Geology 43, 250–280 (1935).

[131] Wang, J. et al. A Molecular Grammar Governing the Driving Forces for Phase Separation of Prion-like RNA Binding Proteins. Cell 174, 688–699 (2018).

[132] Berry, J., Brangwynne, C. P. & Haataja, M. Physical principles of intracellular organization via active and passive phase transitions. Reports on Progress in Physics 81, 046601 (2018).

[133] Boeynaems, S. et al. Spontaneous driving forces give rise to protein- rna condensates with coexisting phases and complex material properties. Proceedings of the National Academy of Sciences 116, 7889–7898 (2019).

[134] Sanchez-Burgos, I., Espinosa, J. R., Joseph, J. A. & Collepardo-Guevara, R. Rna length has a non-trivial effect in the stability of biomolecular condensates formed by rna-binding proteins. PLoS computational biology 18, e1009810 (2022).

[135] Qamar, S. et al. Fus phase separation is modulated by a molecular chaperone and methylation of arginine cation-π interactions. Cell 173, 720–734 (2018).

[136] Murthy, A. C. et al. Molecular interactions underlying liquid-liquid phase separation of the fus low-complexity domain. Nature structural & molecular biology 26, 637–648 (2019).

[137] Rhoads, S. N., Monahan, Z. T., Yee, D. S. & Shewmaker, F. P. The role of post-translational modifications on prion-like aggregation and liquid-phase separation of fus. International journal of molecular sciences 19, 886 (2018).

[138] Tsoi, P. S. et al. Electrostatic modulation of hnrnpa1 low-complexity domain liquid–liquid phase separation and aggregation. Protein Science 30, 1408–1417 (2021).

[139] Li, H.-R., Chiang, W.-C., Chou, P.-C., Wang, W.-J. & Huang, J.-r. Tar dna-binding protein 43 (tdp-43) liquid–liquid phase separation is mediated by just a few aromatic residues. Journal of Biological Chemistry 293, 6090–6098 (2018).

[140] Zacco, E. et al. Rna as a key factor in driving or preventing self-assembly of the tar dna-binding protein 43. Journal of molecular biology 431, 1671–1688 (2019).

[141] McGurk, L. et al. Poly (adp-ribose) prevents pathological phase separation of tdp-43 by promoting liquid demixing and stress granule localization. Molecular cell 71, 703–717 (2018).

[142] Wang, J. et al. A molecular grammar governing the driving forces for phase separation of prion-like rna binding proteins. Cell 174, 688–699 (2018).

[143] Alberti, S. Phase separation in biology. Current Biology 27, R1097–R1102 (2017).

[144] Yang, P. et al. G3bp1 is a tunable switch that triggers phase separation to assemble stress granules. Cell 181, 325–345 (2020).

[145] Schmidt, H. B. & Görlich, D. Nup98 fg domains from diverse species spontaneously phase-separate into particles with nuclear pore-like permselectivity. Elife 4, e04251 (2015).

[146] Linsenmeier, M. et al. Dynamic arrest and aging of biomolecular condensates are modulated by low-complexity domains, RNA and biochemical activity. Nature Communications 13, 1–13 (2022).

