## Supplementary Material for "Protein structural transitions critically transform the network connectivity and viscoelasticity of RNA-binding protein condensates but RNA can prevent it"

Andres R. Tejedor[1,2,‡], Ignacio Sanchez-Burgos[2,‡], Maria Estevez Espinosa[2,3], Adiran Garaizar[2], Rosana Collepardo-Guevara[4,5], Jorge Ramirez[1,*] and Jorge R. Espinosa[2,*]
*[1] Department of Chemical Engineering, Universidad Politécnica de Madrid, José Gutiérrez Abascal 2, 28006, Madrid, Spain.*
*[2] Maxwell Centre, Cavendish Laboratory, Department of Physics, University of Cambridge, J J Thomson Avenue, Cambridge CB3 0HE, United Kingdom.*
*[3] University College London, Gower Street, London WC1E 6BT, United Kingdom*
*[4] Yusuf Hamied Department of Chemistry, University of Cambridge, Lensfield Road, Cambridge CB2 1EW, United Kingdom*
*[5] Department of Genetics, University of Cambridge, Cambridge, CB2 3EH*
*‡ = These authors contributed equally to this work, * = To whom correspondence should be sent.*
(Dated: September 14, 2022)

### SUPPLEMENTARY SECTION I. MODELS AND SIMULATION DETAILS

#### A. HPS-Cation-$\pi$ Model

In order to simulate the different studied proteins and nucleic acids, we employ the recent reparameterization by Das *et al.* [1] of the chemically-accurate coarse-grained (CG) HPS protein model proposed by Dignon *et al.* [2] implemented in the Molecular Dynamics LAMMPS package [3]. For RNA, we use the HPS-compatible CG model proposed by Regy *et al.* [4]. The coarse-grained model resolution, both for proteins and RNA, is of one bead per amino acid and nucleotide. In the model, the intrinsically disordered regions (IDRs) of the proteins are considered as fully flexible polymers, while the structured globular domains are treated as rigid bodies (where their conformations are taken from the Protein Data Bank (PDB) crystalline structure – see SUPPLEMENTARY SECTION I C for the PBD codes) by using the rigid body integrator of LAMMPS [3]. Moreover, the interactions of the structured globular domains are scaled down by a 30% to account for the 'buried' amino acids as proposed by Krainer *et al.* [5]. Also, in this model RNA strands are treated as flexible polymers.

The potential energy of the coarse-grained HPS-Cation-$\pi$ force field is given by:

$$E = E_{\text{Bonds}} + E_{\text{Electrostatic}} + E_{\text{Hydrophobic}} + E_{\text{Cation}-\pi}, \tag{S1}$$

where $E_{\text{Hydrophobic}}$, $E_{\text{Cation}-\pi}$ and $E_{\text{Electrostatic}}$ interactions are only applied between non-bonded beads and $E_{\text{Bonds}}$ between subsequent beads directly bonded to each other.

Bonded interactions between subsequent amino acid protein beads or consecutive RNA nucleotides are described by an harmonic potential:

$$E_{\text{Bonds}} = \sum_{\text{Protein/RNA bonds}} k(r - r_0)^2, \tag{S2}$$

where the equilibrium bond length is $r_0 = 5.0$Å between subsequent nucleotides and $r_0 = 3.81$Å between bonded amino acid beads. The spring constant is $k = 10\,\text{kJ}/(\text{mol}\text{Å}^2)$. The electrostatic interactions, $E_{\text{Electrostatic}}$, among charged amino acids and RNA nucleotides are described by a Coulomb/Debye-Hückel potential of the form:

$$E_{\text{Electrostatic}} = \sum_i \sum_{j<i} \frac{1}{4\pi D} \frac{q_i q_j}{r} e^{-r/\kappa}, \tag{S3}$$

where $q_i$ and $q_j$ represent the charges of the beads $i$ and $j$ (amino acids or nucleotides), $D = 80\epsilon_0$ is the relative dielectric constant of water (being $\epsilon_0$ the electric constant), $r$ is the distance between the $i$th and $j$th beads, and $\kappa$= 1 nm is the Debye screening length that mimics the implicit solvent (water and ions) at physiological salt concentration ($\sim$150mM of NaCl) [2].

The hydrophobic interactions between different amino acid types and nucleotides are built upon a scale of amino acid and RNA nucleotide hydrophobicity based on a statistical potential derivation from contacts in PDB structures, and implemented through the functional form of an Ashbaugh/Hatch potential (for further details see References [2, 4, 6, 7]):

$$E_{\text{Hydrophobic}} = \sum_i \sum_{j<i} \begin{cases} 4\epsilon_{ij} \left[ \left(\frac{\sigma_{ij}}{r}\right)^{12} - \left(\frac{\sigma_{ij}}{r}\right)^{6} \right] + (1-\lambda_{ij})\epsilon_{ij}, & r < 2^{1/6}\sigma_{ij} \\ \lambda_{ij} 4\epsilon_{ij} \left[ \left(\frac{\sigma_{ij}}{r}\right)^{12} - \left(\frac{\sigma_{ij}}{r}\right)^{6} \right], & \text{otherwise}, \end{cases} \tag{S4}$$

where $\lambda_i$ and $\lambda_j$ are parameters that account for the hydrophobicity of the $i$th and $j$th interacting particles respectively, being $\lambda_{ij} = (\lambda_i + \lambda_j)/2$. The excluded volume of the different residues/nucleotides is given by $\sigma_i$ and $\sigma_j$, where $\sigma_{ij} = (\sigma_i + \sigma_j)/2$, and $r$ is the distance between the $ij$ particles. $\epsilon_{ij}$ (0.2 kcal/mol) is a fitting parameter to reproduce the experimental single-IDR radius of gyration ($R_g$) [2]. When at least one of the $ij$ amino acids is part of a structured globular domain, $\lambda_{ij}$ is scaled by a factor of 0.7 to account for the 'buried' amino acids in globular domains [5]. The specific values for each amino acid and nucleotide $\sigma$, $q$, and $\lambda$ parameters can be found in References: Dignon *et al.* for proteins [2] and Regy *et al.* for RNA [4].

Finally, we consider an extra term for describing cation-$\pi$ (c-$\pi$) interactions (only for the following set of pairs of amino acids (c-$\pi$:{Arg-Phe, Arg-Trp, Arg-Tyr, Lys-Phe, Lys-Trp and Lys-Tyr}):

$$E_{\text{cation-}\pi} = \sum_{i \in \{\text{c-}\pi\}} \sum_{\substack{j \in \{\text{c-}\pi\} \\ j<i}} 4\epsilon_{ij} \left[ \left(\frac{\sigma_{ij}}{r}\right)^{12} - \left(\frac{\sigma_{ij}}{r}\right)^{6} \right], \tag{S5}$$

where $\sigma_{ij}$ is the same as in the hydrophobic interactions and $\epsilon_{ij}$ is $\epsilon_{ij} = 3.0 \text{kcal mol}^{-1}$ for all six cation-$\pi$ pairs as proposed in Ref. [1] (Approach 1). Consistently, the interaction of these amino acids is scaled down by a 30% when they are found in structured globular domains.

#### B. Mpipi model

Furthermore, we also report results in Fig. 5 of the main text employing the Mpipi force field [8]. This model describes almost quantitatively the temperature-dependent LLPS phase behaviour of different protein condensates such as those of FUS or hnRNPA1. In addition, this model correctly predicts the multiphase behaviour of PolyR/PolyK/PolyU systems, and recapitulates experimental LLPS trends for sequence mutations on FUS, DDX4 NTD, and LAF-1 RRG domain variants [8]. Within this force field, electrostatic interactions are modelled with a Coulomb/Debye-Huckel term as in the HPS model (Eq. S3), being $\kappa^{-1} = 795$ pm, $\epsilon_r = 80$ and the charges -0.75$e$ for the different nucleotides: A, C, G, U; and +0.75$e$ for amino acids such as R and K, +0.375$e$ for H, and -0.75$e$ for D and E residues. The cut-off for the electrostatic interactions is set at 3.5 nm (see Ref. [8] for further details on the electrostatic term of the model). Non-bonded interactions between protein/RNA beads are modelled via the Wang–Frenkel potential [9]:

$$E_{WF} = \sum_{i,j} \epsilon_{ij} \alpha_{ij} \left[ \left( \frac{\sigma_{ij}}{r_{ij}} \right)^{2\mu_{ij}} - 1 \right] \left[ \left( \frac{R_{ij}}{r_{ij}} \right)^{2\mu_{ij}} - 1 \right]^{2\nu_{ij}} \tag{S6}$$

where

$$\alpha_{ij} = 2\nu_{ij} \left( \frac{R_{ij}}{\sigma_{ij}} \right)^{2\mu_{ij}} \left[ \frac{2\nu_{ij} + 1}{2\nu_{ij} \left[ \left( \frac{R_{ij}}{\sigma_{ij}} \right)^{2\mu_{ij}} - 1 \right]} \right]^{2\nu_{ij}+1} \tag{S7}$$

representing $\sigma$ the molecular diameter of each residue/nucleotide and $\epsilon$ the interaction strength between distinct amino acids and nucleotides ($i$ and $j$). $\nu_{ij}$ and $R_{ij}$ are constant model parameters set to $\nu_{ij} = 1$ and $R_{ij} = 3\sigma_{ij}$ for every interaction, while $\sigma_{ij}$, $\epsilon_{ij}$ and $\mu_{ij}$ are specified for each pair of interaction in Ref. [8]. Finally, in this model bond energy is computed with an harmonic bond potential of the following form:

$$E_{bond} = \sum_b \frac{1}{2} k(r_b - r_0) \tag{S8}$$

where $b$ is the total number of bonds, $r_b$ is the bond distance, $k$=8.03 Jmol$^{-1}$pm$^{-2}$ is the spring constant and $r_0$ is the bond reference position, set to 381 pm and 500 pm for protein and RNA bonds respectively. For further details on this force field and the full list of the model parameters please see Ref. [8].

### C. Protein sequences and PDB of the structured domains

#### FUS

MASNDYTQQATQSYGAYPTQPGQGYSQQSSQPYGQQSYSGYSQSTDTSGYGQSSYSSYGQSQNTGYGTQSTPQGYGSTGGYGSS QSSQSSYGQQSSYPGYGQQPAPSSTSGSYGSSSQSSSYGQPQSGSYSQQPSYGGQQQSYGQQQSYNPPQGYGQQNQYNSSSGGGG GGGGGGNYGQDQSSMSSGGGSGGGYGNQDQSGGGGSGGYGQQDRGGRGRGGSGGGGGGGGGGYNRSSGGYEPRGRGGGRG GRGGMGGSDRGGFNKFGGPRDQGSRHDSEQDNSDNNTIFVQGLGENVTIESVADYFKQIGIIKTNKKTGQPMINLYTDRETGKL KGEATVSFDDPPSAKAAIDWFDGKEFSGNPIKVSFATRRADFNRGGGNGRGGRGRGGPMGRGGYGGGGSGGGGRGGFPSGGG GGGGQQRAGDWKCPNPTCENMNFSWRNECNQCKAPKPDGPGGGPGGSHMGGNYGDDRRGGRGGYDRGGYRGRGGDRGGF RGGRGGGDRGGFGPGKMDSRGEHRQDRRERPY

The following Protein Data Bank (PDB) codes were used to build the globular structured domains of FUS (residues from 285–371 (PDB code: 2LCW) and from 422–453 (PDB code: 6G99)).

#### FUS-PLD

MASNDYTQQATQSYGAYPTQPGQGYSQQSSQPYGQQSYSGYSQSTDTSGYGQSSYSSYGQSQNTGYGTQSTPQGYGSTGGYGSS QSSQSSYGQQSSYPGYGQQPAPSSTSGSYGSSSQSSSYGQPQSGSYSQQPSYGGQQQSYGQQQSYNPPQGYGQQNQYNS

#### A-LCD-hnRNPA1

MASASSSQRGRSGSGNFGGGRGGGFGGNDNFGRGGNFSGRGGFGGSRGGGGYGGSGDGYNGFGNDGSNFGGGGSYNDFGNYN NQSSNFGPMKGGNFGGRSSGPYGGGGQYFAKPRNQGGYGGSSSSSSYGSGRRF

### D. Simulation details

#### Potential of mean force calculations in all-atom simulations

We estimate the potential of mean force (PMF) by performing a set of atomistic Umbrella Sampling MD simulations for the GYNGFG segment (PDB code: 6BXX) within the A-LCD-hnRNPA1 protein. The simulations are performed with explicit water and ions using the a99SB-*disp* force field for protein, water and ions [10] using the GROMACS 2018 package [11]. The starting configurations for each simulations consisted of four stacked peptides (two pairs of peptides, stacked on top of each other forming a ladder) taken from cross-$\beta$-fibril structures resolved crystallographycally [12]. Acetyl and N-methyl capping groups were added to the termini of each peptide. For each 4-peptide system, we performed two sets of simulations. First, we compute the interactions among the four peptides when they remain in the cross-$\beta$-fibril structure by imposing positional restraints on all the heavy atoms (1000 kJ mol$^{-1}$ nm$^{-2}$; yellow curve in Figure 1a of the main text) in all directions (except for the dissociating peptide in the pulling direction); this ensures that the four individual peptides maintain their secondary structure during the simulation. In the second set of simulations, for the interactions among disordered peptides, we allow all peptides to freely sample their configurational landscape (purple curve in Figure 1a of the main text). In both cases, we kept the positions of the central C$\alpha$ atoms fixed to control the relative distances between peptides (except in the direction of the reaction coordinate for the dissociating peptide). For the visualization of the PMF atomistic simulations, we have utilized VMD software (version 1.9.1) [13].

As the reaction coordinate, we use the distance between the center-of-mass (COM) of the 'dissociating' peptide and the closest peptide to it along the pulling direction. In the case of the disordered peptides, rather than the COM distance, we took the distance between the central C$\alpha$'s in the aforementioned chains, to avoid noise due to changes in the COM from fluctuations in the peptide conformation. We spaced umbrella windows approximately every 0.2Å along the reaction coordinate, from 4 to 30Å (where longest range interactions completely vanish). To sample the steep potential among rigid structures, a spring constant of 24000 kJ mol$^{-1}$ nm$^{-2}$ is used.

For the ordered peptides, we ran simulations of about 25 ns per umbrella window. For the disordered peptides, a simulation time of 100 ns is necessary. Two independent simulation trajectories were performed for each Umbrella Sampling PMF window (Fig. 1a of the main text). The employed timescales guarantees uncertainties of 1-1.5$k_BT$ at most in the dissociation profiles of the disordered LARKS. For the integration of the equations of motion, we used the Verlet algorithm with a time step of 1 fs. To constrain bond lengths and angles, we used the LINCS algorithm [14] with an order of 8 and 2 iterations. The cut-off for the Coulomb and van der Waals interactions was chosen at a conservative value of 1.4 nm. For electrostatics, we used Particle Mesh Ewald (PME) [15] of 4th order with a Fourier spacing of 0.12 nm and an Ewald tolerance of $1.5{\cdot}10^{-5}$. The simulations were performed in the NpT ensemble. The temperature and pressure were kept constant using a Nosé–Hoover thermostat [16] at T = 300 K (with 1 ps relaxation time) and a Parrinello–Rahman [17] isotropic barostat at p = 1 bar (with a 1 ps relaxation time), respectively. A NaCl concentration of 0.1 M was used throughout.

Each simulation system comprises of a box of approximately 6 x 6 x 8 nm$^3$. We used $\sim$ 7500 water molecules. All our systems were electroneutral (i.e., the total net charge was zero). After solvation of the peptides, we perform an energy minimization with a force tolerance of 1000 kJ mol$^{-1}$ nm$^{-1}$, followed by a short, 1000 ps NpT equilibration both with positional restraints (10000 kJ mol$^{-1}$ nm$^{-2}$) for the heavy atoms of the chains in the three directions of space. The analysis of the simulations was carried out using the WHAM [18] analysis implemented in GROMACS (version 2018.2-foss-2018b). The first 10% of the simulations was discarded as equilibration time.

**Residue-resolution coarse-grained simulations**

Within our residue-resolution simulations (Mpipi and HPS-Cation-$\pi$ models), we employ Direct Coexistence (DC) simulations [19] to compute phase diagrams in the NVT ensemble (i.e. constant number of particles (N), volume (V) and temperature (T)), for which we use a Langevin thermostat [20] with a relaxation time of 50 ps for both the HPS-Cation-$\pi$ and Mpipi models. Since all our potentials are continuous and differentiable, we perform all our simulations using the LAMMPS Molecular Dynamics package (version 21st of July of 2020) [21]. Simulations of full-FUS containing globular domains are performed using the LAMMPS rigid body Nosé–Hoover thermostat [22–24] in combination with a Langevin thermostat [20]. Periodic boundary conditions are used in the three directions of space. The timestep chosen for the Verlet integration of the equations of motion is 10 fs for both models. Disorder-to-order transitions implemented through our dynamical model in bulk phases are studied running simulations in the NpT ensemble at p=0 bar using a Nosé-Hover barostat [3] and thermostat [16] with relaxation times of 50 ps and 5 ps respectively, and integration time steps of 10 fs. We run independent seeds for FUS and A-LCD-hnRNPA1 in order to increase the statistical significance of our measurements, specially in RNA-protein mixtures for all the studied range of concentrations (including also the cases in absence of RNA). More precisely, we run 3 different trajectories of A-LCD-hnRNPA1-polyU mixtures (Fig. 5a in the main text) and 5 independent runs of FUS-polyU condensates (Fig. 4c in the main text) at every polyU/protein mass ratio. The results we show in the main text are obtained by averaging the number of disorder-to-order transitions using the different independent trajectories. To compute viscoelastic properties and droplet intermolecular contact frequencies, we run NVT simulations using a cubic simulation box. The volume of each box is set to the bulk equilibrium density at the corresponding temperature. For viscosity calculations, we perform simulations of about 3$\mu$s. The integration timestep and relaxation time of the thermostat for the viscoelastic properties calculations are the same used in our DC simulations and NpT simulations described above. We have used Ovito (version 3.7.3) [25] for visualization of coarse-grained simulations. The system size for each of the studied protein condensates are: 200 protein replicas for A-LCD-hnRNPA1 simulations, 100 replicas for FUS-PLD simulations, and 48 protein replicas for full-sequence FUS simulations. Simulations for these protein condensates in presence of polyU are performed by adding the corresponding amount of polyU (of 125-nucleotide strands) that reproduces the polyU/protein mass ratios described along the manuscript.

**Potential of mean force calculations with the HPS-Cation-$\pi$ Model**

Similarly to our atomistic simulations, we perform PMF calculations employing the (sequence-dependent coarse-grained) HPS-Cation-$\pi$ model with both the Reference (Disorder) and Ageing (Structured) parametrizations described in the main text and Supplementary Table II. These two parametrizations essentially differ in the interaction strength $\epsilon$ and the angular constant $k_{ang}$. We model a 4-peptide aggregate (sequence $_{58}$GYNGFG$_{63}$) using both parametrizations starting from a close-packed configuration of four (aligned) peptides at its lowest possible interaction energy. From that configuration and using both set of parameters, we gradually dissociate one of the peptides from the

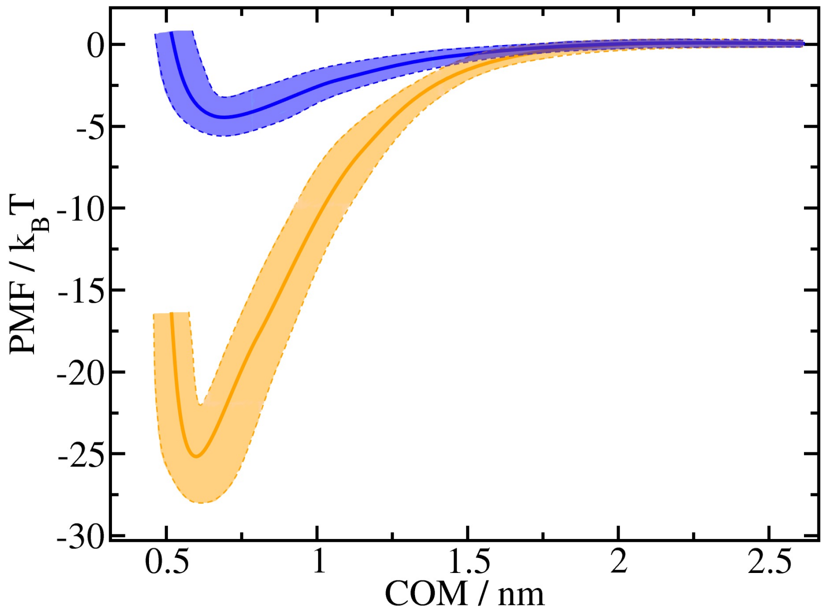


**Supplementary Figure 1:** Potential of Mean Force (PMF) dissociation curve of a 6-amino acid segment ($_{58}$GYNGFG$_{63}$) found in the A-LCD-hnRNPA1 sequence from a $\beta$-sheet structure formed by 4 LARKS (of the same sequence) as a function of the center of mass distance (COM) using the HPS-Cation-$\pi$ force field [1, 2]. PMF simulations are conducted at T/T$_c$=0.95 of the pure A1-LCD-hnRNPA1 system. The blue curve represents the PMF dissociation profile among peptides with a disordered-like interaction (corresponding to the Reference Model: i.e., HPS-Cation-$\pi$ parameters), while the orange curve depicts the dissociation curve among the same segments with structured-like interactions (corresponding to the structured LARKS interaction of the Ageing model; Supplementary Table II). The statistical uncertainty of both PMF calculations (depicted by colour bands) has been obtained by bootstrapping n = 4 independent simulations.

other segments as performed in our atomistic simulations (Fig. 1a of the main text). As the reaction coordinate, we use the distance between the center-of-mass (COM) of the 'dissociating' peptide and the closest peptide to it along the pulling direction. Furthermore, we impose positional restraints (1000 kJ mol$^{-1}$ nm$^{-2}$ ) on all beads except for the dissociating peptide in the pulling direction, while a spring constant of 20000 kJ mol$^{-1}$ nm$^{-2}$ is used to sample each window around a given COM distance. The Umbrella Sampling windows are spaced in 0.3 Å from each other.

Calculations are performed in 200 Å x 200 Å x 200 Å simulation boxes for 40 ns each at T/T$_c$=0.95 of the pure A1-LCD-hnRNPA1 system. We integrate the equations of motion using the Verlet algorithm with a time step of 2 fs. PMF simulations were performed in the NVT ensemble, keeping constant the temperature using a Nosé–Hoover thermostat [16] with 0.5 ps relaxation time. Results from the Reference Model (blue curve) and Ageing Model (orange curve) are represented in Supplementary Figure 1.

### SUPPLEMENTARY SECTION II. OBTAINING PHASE DIAGRAMS VIA DIRECT COEXISTENCE SIMULATIONS

To calculate the coexisting densities of the phase diagrams, we employ the Direct Coexistence method [19, 26, 27]. Within this scheme, the two coexisting phases are simulated by preparing periodically extended slabs of the two phases, the condensed and the diluted phase, in the same simulation box. Once our DC simulations have reached equilibrium, we compute the density profile along the long axis of the box, and thus, we extract the density of the two coexisting phases (i.e., Figs. 1d and 2a of the main text). We note that once equilibrium is reached, it is equivalent to perform several short independent trajectories or a single long simulation as long as the total simulated time is equivalent. To estimate the critical point of the phase diagrams, we use the universal scaling law of coexistence densities near a critical point [28], and the law of rectilinear diameters [29]:

$$\big(\rho_l(T) - \rho_v(T)\big)^{3.06} = d\bigg(1 - \frac{T}{T_c}\bigg) \tag{S9}$$

and

$$(\rho_l(T) + \rho_v(T))/2 = \rho_c + s_2(T_c - T) \tag{S10}$$

where $\rho_l$ and $\rho_v$ refer to the coexisting densities of the condensed and diluted phases respectively, $\rho_c$ is the critical density, $T_c$ is the critical temperature, and $d$ and $s_2$ are fitting parameters. The critical temperatures are provided in the Supplementary Table I for the reference model. Please note that the value of $T_c$ barely changes when the dynamical algorithm is applied. The error bars in the coexisting densities of the phase diagrams shown in the main text (i.e., Figs. 1d, 2a and 4a) have been estimated by applying a block average analysis of the bulk density of each phase across our Direct Coexistence simulations (i.e., discarding regions near the condensate interfaces). On the other hand, the uncertainty in the critical temperature and density is evaluated by considering distinct possible fits of the law of rectilinear diameters and critical exponents using different coexistence densities ranging within the previously computed uncertainty.

| FUS | FUS-PLD | A1-LCD-hnRNPA1 |
|---|---|---|
| $(395 \pm 5)\,K$ | $(350 \pm 5)\,K$ | $(412 \pm 5)\,K$ |

**Supplementary Table I:** Obtained critical temperatures from simulations of the different studied proteins.

### SUPPLEMENTARY SECTION III. AGEING DYNAMIC ALGORITHM

To dynamically mimic the structural diversity of proteins during condensate maturation, we implement a time-dependent and local-dependent algorithm based on our previous work from Ref. [30]. Low-complexity aromatic-rich kinked segments (LARKS) as ($_{58}$GYNGFG$_{63}$ within the A-LCD-hnRNPA1 sequence, and $_{37}$SYSGYS$_{42}$, $_{54}$SYSSYGQS$_{61}$ and $_{77}$STGGYG$_{82}$ within the FUS sequence) can spontaneously transition from their fully disordered state to form inter-peptide structured $\beta$-sheets depending on the local environment. For effectively considering that behaviour in our coarse-grained simulations, every 100 simulation timesteps, our dynamical algorithm evaluates whether the conditions around each fully disordered LARKS are favorable for undergoing an 'effective' disorder-to-order cross-$\beta$-sheet transition, and thus, modifying the protein parameters given in the Supplementary Table II (derived from all-atom simulations; Fig. 1 of the main text), to those corresponding to inter-peptide structured $\beta$-sheet motifs. These conditions are evaluated on the central bead of each LARKS (the specific amino acid for which the coordination calculations are performed is detailed in Supplementary Table II). We make use of a distance criterion, so that if 4 peptides are within a certain cut-off distance ($r_{cut}$ in Supplementary Table II), the average effective interaction in the aforementioned domains is substituted by that obtained from all-atom PMF calculations for the residues belonging to such sequence (e.g., $\lambda$ of the corresponding amino acids is updated to a global $\lambda_{ordered}$ from Supplementary Table II for the whole LARKS once the transition has taken place). Once a disorder-to-$\beta$-sheet transition takes place, it may be reversible in case one of the proteins due to thermal fluctuations may leave the structured inter-peptide domain beyond a certain distance ($r_{cut,reverse}$ in Supplementary Table II). Since the identity of the amino acids involved in protein structural changes is virtually modulated upon the transition, we also assign an average mass to the new structured LARKS residues that keeps constant the total mass of the protein, and hence, the mass of the amino acids forming structured cross-$\beta$-sheet domains (values reported in Supplementary Table II as $m_{ordered}$). Additionally, to account for the higher rigidity of inter-peptide $\beta$-sheet motifs upon the structural transition, we introduce an angular term to the total energy (equation S1) that follows the next equation:

$$E_{\text{Angles}} = \sum_{\text{Angles}} k_{ang}(\theta - \theta_0)^2, \tag{S11}$$

where we set $\theta_0 = 180°$ and $k_{ang} = 5\ kcal\ mol^{-1}\ rad^{-2}$. To carry out these simulations, we used the USER-REACTION [31] package of LAMMPS (version 21st of July of 2020) which allows to change the topology of the underlying system components on a time-dependent and local-dependent manner.

| | A-LCD-hnRNPA1 | FUS (1) | FUS (2) | FUS(3) |
|---|---|---|---|---|
| Central amino acid | $N_{60}$ | $S_{39}$ | $S_{57}$ | $G_{79}$ |
| $m_{ordered}$ / (g mol$^{-1}$) | 99.27 | 107.44 | 107.48 | 86.92 |
| $\lambda_{ordered}$ / (kcal mol$^{-1}$) | 3.2 | 2.8 | 3.5 | 2.2 |
| $\sigma_{ordered}$ / Å | 5.33 | 5.52 | 5.52 | 5.13 |
| $r_{cut}$ / Å | 8 | 7 | 7 | 7.75 |
| $r_{cut,reverse}$ / Å | 20 | - | - | 20 |

**Supplementary Table II:** Set of parameters employed for residues belonging to structured inter-peptide $\beta$-sheet motifs in HPS-Cation-$\pi$ simulations. Note that FUS (1), FUS (2) and FUS (3) correspond to $_{37}$SYSGYS$_{42}$, $_{54}$SYSSYGQS$_{61}$ and $_{77}$STGGYG$_{82}$ sequences respectively. Moreover, since the binding interaction strength of FUS(1) and FUS(2) LARKS cross-$\beta$-sheet fibrils is high enough to (almost always) prevent dissociation events, a backward reaction for those two LARKS has not been set in order to accelerate the performance of the dynamical ageing algorithm. Please note that an analogous reparametrization has been performed for the Mpipi simulations shown in Fig. 5b of the main text.

The criterion that at least four peptides should be in close contact to trigger a disorder-to-order transition has been chosen based on the following arguments. Four interacting peptides is the minimal system where all the different types of stacking and hydrogen bonding interactions that stabilize the $\beta$-sheet fibrillar ladder are fulfilled; i.e., two interacting steps of the ladder each made of a pair of $\beta$-sheet peptides. Thus, considering fewer interacting peptides (e.g. only three or two) would severely underestimate the strength of interactions among ordered LARKS in our atomistic simulations, and subsequently, erroneously preclude the formation of kinetically arrested states at the coarse-grained level. Our atomistic simulations (Fig. 1 of the main text) and those from our previous work (Ref. [30]), show that the strength of interactions among four ordered peptides is already high enough for the peptides to remain stably bound upon thermal fluctuations. Hence, if we made the criterion even more stringent (i.e., by requiring clustering of five or more peptides), the strength of interactions among the system would remain consistent with kinetic arrest at the coarse-grained level. However, a more stringent criterion would now render the coarse-grained simulations prohibitive expensive. That is, much longer simulation timescales would be needed to capture the rarer higher-density fluctuations that could result into the spontaneous formation of clusters of five or more peptides as opposed to the more frequent fluctuations that yield clusters of four peptides (as considered within our dynamical algorithm presented here).

To demonstrate that condensate ageing (induced by disorder-to-order transitions) cannot take place in our model unless it is mediated by strong density fluctuations characteristic of a phase-separated droplet, we perform simulations of full-FUS using the dynamical ageing model at the critical temperature and density obtained in Fig. 2a. As it can be seen in the Supplementary Figure 2, the percentage of cross-$\beta$-sheet transitions over time is roughly zero. Since the system at the critical conditions cannot undergo LLPS, and as consequence display strong (high/low) density fluctuations, the emergence of cross-$\beta$-sheet transitions over time is extremely scarce and quickly dissociate once (if) they are formed.

### SUPPLEMENTARY SECTION IV. ANALYSIS OF THE NETWORK BY A PRIMITIVE PATH ANALYSIS

The primitive path analysis (PPA) [32] has been extensively used in polymer physics simulations to investigate the underlying network structure of entangled polymer melts. The algorithm minimizes the contour length of all chains in a system, while keeping the end monomers fixed and respecting the network topology by not allowing chains to cross each other, and allows to obtain different properties that characterize the rubber elastic properties of entangled polymers (such as the number of entanglements, or the tube diameter [33]). In this work, we employ a modified version of the analysis proposed by Sukumaran *et al.* [32] to study the connectivity of the condensed-phase network along condensate maturation. We carry out long NpT simulations to allow the number of cross-$\beta$-sheet transitions in each of the different studied systems to reach a steady state. Then, we apply PPA to the resulting configuration to obtain information about the concentration of elastically active strands in the system. In our modified version of the algorithm, instead of fixing the positions of the molecular ends in space, we freeze the coordinates of the inter-petide $\beta$-sheet structures. Then, PPA proceeds as an energy minimization in which the intramolecular excluded volume interactions have been switched off, and the equilibrium bond length has been set to zero, so that the contour length of the strands between the transitioned structured LARKS domains are minimized while preserving the underlying network connectivity. At the end of the minimization, we are left with the minimal network of elastically active protein strands that contribute to the formation of a rubbery plateau in $G(t)$, as shown in Fig. 3c of the main text. If the network percolates $G(t)$ shows a clear plateau, whereas if the proteins form disconnected clusters, $G(t)$ will decay to zero with a slightly higher viscosity. We employ a modified version of the LAMMPS Molecular

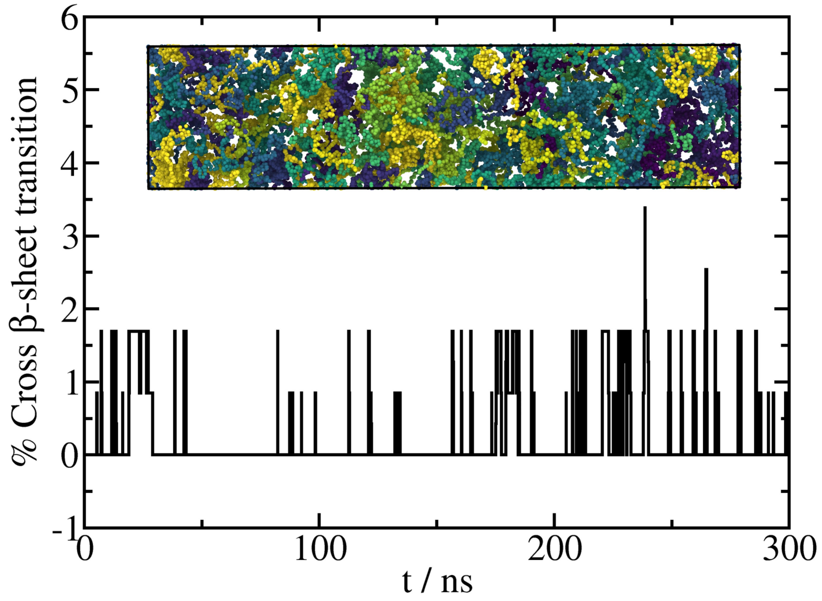


**Supplementary Figure 2:** Number of inter-peptide $\beta$-sheet transitions as a function of time for a full-FUS system at the critical temperature and density obtained in Fig. 2a of the main text. As it can be seen, the rate of structural transitions is extremely low, and when sporadic cross-$\beta$-sheet transitions take place, inter-peptide $\beta$-sheet clusters quickly dissociate due to strong low-density fluctuations. A snapshot of the system has been also included in which each FUS replica is depicted by a different colour.

Dynamics simulation package (version 21st of July of 2020) [3] to perform the primitive-path analysis, running energy minimization simulations using 1e5 iterations with a tolerance of 1.0e-6 for both the total force and the energy. In movies SM1 and SM2 we show examples of how the PPA proceeds for the two systems whose underlying networks are depicted in Fig 3c of the main text.

### SUPPLEMENTARY SECTION V. TIME-EVOLUTION OF CROSS-$\beta$-SHEET TRANSITIONS THROUGH A SECOND-ORDER REACTION ANALYSIS.

We have employed a second-order reaction analysis to study the time-evolution of the number of disorder-to-$\beta$-sheet transitions within both A-LCD-hnRNPA1 and FUS condensates as a function of RNA concentration using 125-nt polyU chains (see Figs. 4c and 5a-b in the main text) and at constant temperature (T=0.97T$_c$ for A-LCD-hnRNPA1 and T=0.96T$_{c,FUS}$ for FUS condensates). We use the same reaction mechanism for both systems, despite the evolution of A-LCD-hnRNPA1 transitions are driven by only one reacting site whereas the curves provided in Fig. 4c are obtained by averaging the three LARKS motifs of FUS.

We define $[A]$ as the concentration of reactive sites in their disordered state and $[B]$ as the concentration of structured domains. We assume that the A and B sites react according to the following mechanism:

$$\begin{aligned} 2A &\xrightarrow{k_{dir}} 2B \\ A + B &\xrightarrow{k_{dir}} 2B \\ A &\xleftarrow{k_{inv}} B, \end{aligned} \tag{S12}$$

where $k_{dir}$ is the kinetic constant for the direct reaction and $k_{inv}$ is the kinetic constant for the inverse reaction. Defining the equilibrium constant as $K_{eq} = \frac{k_{dir}}{k_{inv}}$ and assuming perfect mixing of all the species in the system, we can write the following system of ordinary differential equations for the evolution of the concentrations of $A$ and $B$

$$\begin{aligned}\frac{d[A]}{dt} &= -2k_{dir}[A]^2 - k_{dir}[A][B] + \frac{k_{dir}}{K_{eq}}[B] \\ \frac{d[B]}{dt} &= 2k_{dir}[A]^2 + k_{dir}[A][B] - \frac{k_{dir}}{K_{eq}}[B].\end{aligned} \tag{S13}$$

This system of equations can be easily integrated using an adaptive time step algorithm for different values of the parameters $k_{dir}$ and $K_{eq}$, and used to the time-evolution of cross-$\beta$-sheet transitions, so the kinetic constants are obtained as fitting parameters. We performed the analysis with our own Python code implemented in RepTate (version 1.1.1 20200602) [34]. The value for the direct reaction constant $k_{dir}$ is shown in the main text for both systems, and here we provide the optimal values of both constants in Suppelemtary Table III for A-LCD-hnRNPA1/polyU mixtures and Supplementary Table IV for FUS/polyU mixtures. Please note that the error in the estimates of $K_{eq}$ may be specially significant in the case of A-LCD-hnRNPA1 ageing, because (1) A-LCD-hnRNPA1/polyU transitions are mostly irreversible under some of the studied conditions (i.e., at low RNA concentration), and (2) the steady state is not well-captured in some cases possibly yielding misleading values of the equilibrium constant $K_{eq}$. By running longer simulations, better estimates of the parameters can be easily obtained. We have performed 5 independent trajectories (by changing the initial particle velocity) for the simulations shown in Figs. 1e, 1f, 4c, 5a and 5b of the main text.

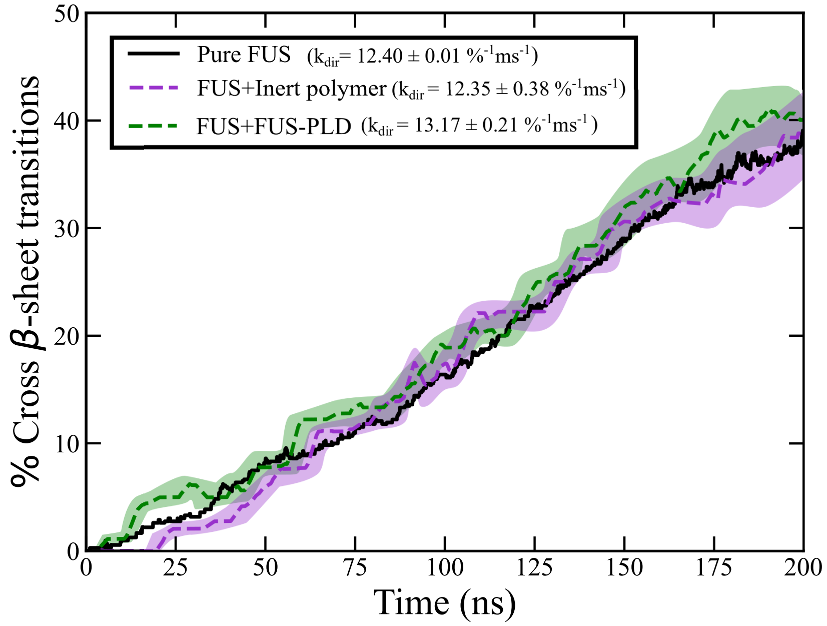


**Supplementary Figure 3:** Time-evolution of inter-protein $\beta$-sheet transitions (in percentage) within bulk condensates at $T/T_{c,FUS}$=0.96 for pure full-FUS (black solid line), a mixture of full-FUS and inert polymers (purple dashed line), and a mixture of full-FUS with FUS-PLD (green dashed line). While full-FUS is the main component, the monomeric concentration of inert polymers and FUS-PLD replicas has been set to be equivalent to the monomeric concentration of the 0.24 polyU/FUS (mass ratio) condensate shown in Fig. 4c of the main text. Shading bands depict the statistical uncertainty of the green and purple curves, which was obtained by bootstrapping n = 2 independent trajectories, whereas that of the black curve was computed using n = 5 independent simulations.

### SUPPLEMENTARY SECTION VI. LOCAL ORDER AND SPHERICITY PARAMETERS

To determine the sphericity ($\Phi$) of the condensates, as shown in Fig. 3d of the main text, we first need to identify the protein replicas belonging to the phase-separated droplet. To do so, we initially determine the average number of neighbours that each amino acid of a given protein replica presents. By setting a threshold cut-off distance of 7Å, we compute the histogram of neighbours (Supplementary Figure 4) for both condensed and diluted phases in bulk. The number of neighbours that minimize the mislabeling of proteins is 0.2 (Supplementary Figure 4). Now only taking

| A-LCD-hnRNPA1 (HPS-Cation-$\pi$ model) | | | A-LCD-hnRNPA1 (Mpipi model) | | |
|---|---|---|---|---|---|
| RNA/protein mass ratio | $K_{eq}$ | $k_{dir}$ ($\%^{-1}ms^{-1}$) | RNA/protein mass ratio | $K_{eq}$ | $k_{dir}$ ($\%^{-1}ms^{-1}$) |
| 0.00 | 1.352 ± 0.015 | 14.480 ± 0.035 | 0.00 | 6.6e+06 ± 1.6e+01 | 12.520 ± 0.010 |
| 0.059 | 2.5e+05 ± 1.4e+05 | 15.480 ± 0.015 | 0.117 | 3.0e+07 ± 3.3e+02 | 11.930 ± 0.016 |
| 0.088 | 4.2e+06 ± 3.7e+01 | 13.930 ± 0.021 | 0.235 | 7.6e+07 ± 7.4e+01 | 11.190 ± 0.015 |
| 0.117 | 1.9e+07 ± 1.9e+02 | 13.110 ± 0.010 | 0.293 | 5.7e+07 ± 1.1e+02 | 8.5310 ± 0.0054 |
| 0.176 | 1.7e+07 ± 1.0e+02 | 7.689 ± 0.014 | | | |

**Supplementary Table III:** Kinetic constants obtained from fits to a second-order reaction model for A-LCD-hnRNPA1/polyU bulk condensates using both HPS-Cation$-\pi$ and Mpipi force fields and at different polyU/protein mass ratios.

| FUS (HPS-Cation-$\pi$ model) | | |
|---|---|---|
| RNA/protein mass ratio | $K_{eq}$ | $k_{dir}$ ($\%^{-1}ms^{-1}$) |
| 0.000 | 7.3e+04 ± 7.7e+01 | 12.409 ± 0.019 |
| 0.118 | 1.5e+05 ± 1.9e+02 | 12.371 ± 0.020 |
| 0.236 | 2.1 ± 0.054 | 11.532 ± 0.023 |

**Supplementary Table IV:** Kinetic constants obtained from fits to a second-order reaction model for FUS/polyU bulk condensates using the HPS-Cation$-\pi$ model at different polyU/FUS mass ratios.

into account the condensed phase, we identify the biggest droplet by considering only proteins belonging to the same cluster (as long as they are connected with other proteins in the same condensate within a cut-off distance lower than 7Å).

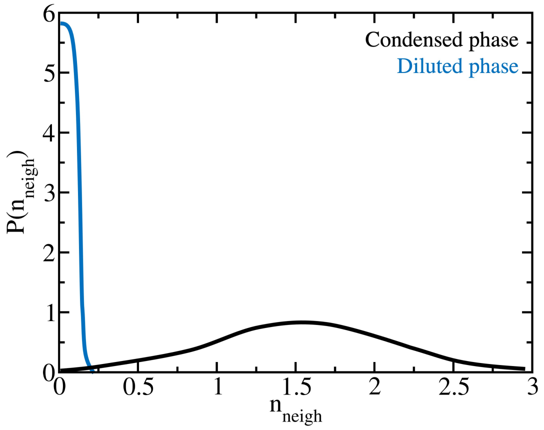


**Supplementary Figure 4:** Histogram of the number of neighbours that residues of FUS-PLD proteins display within the condensed and the dilute phase in bulk conditions at T/T$_{c,FUS}$=0.81. Two amino acids are considered neighbours if they are found within a cut-off distance of 7Å.

Then, by only focusing on the main droplet, we perform a Solvent Available Surface Area (SASA) analysis [35] that computes the area of the condensate that is available for a solvent spherical particle of radius 3Å (i.e., roughly of the same size than a water molecule) as well as the volume enclosed in such geometric shape. For such purpose, we make use of the *gmx sasa* analysis tool in the GROMACS (version 2018.2-foss-2018b) software package [11]. Lastly, the estimate of the condensate sphericity is computed by using the following expression [36]:

$$\Phi(t) = \frac{\pi^{1/3} 6^{2/3} V(t)^{2/3}}{A(t)} \tag{S14}$$

where $V(t)$ and $A(t)$ refer to the volume and area of the droplet for a given time $t$. This parameter is defined within an interval [0,1], taking the value of 1 for a perfect sphere.

### SUPPLEMENTARY SECTION VII. CALCULATION OF MOLECULAR CONTACT PROBABILITIES

Landscapes of intermolecular protein contact probabilities are computed from NVT simulations at constant temperature and equilibrium bulk density. These results are obtained from the same simulations employed to calculate the viscoelastic properties within the protein condensates (see SUPPLEMENTARY SECTION I D and SUPPLEMENTARY SECTION VIII) for both the reference and the ageing model. We use our own Python (version 3.7.2-GCCcore-8.2.0) code to perform this analysis. We can approximately estimate the variation in binding free energy along droplet maturation utilizing the intermolecular maps of contact probability obtained from the NVT trajectories that contain ∼100 independent configurations. We employ the 'smart' cut-off analysis introduced in our previous work [37]. Within that scheme, the cut-off distance is set to $1.2\sigma_{ij}$, where $\sigma_{ij}$ accounts for the mean excluded volume of each specific pair $i$th and $j$th amino acids. Since the minimum of the pairwise potential (Eq. (S5)) is located at $2^{1/6}\sigma_{ij} \approx 1.122\sigma_{ij}$, we set our cut-off distance slightly beyond that point, at $1.2\sigma_{ij}$, to ensure significant attractive binding.

The intermolecular contact map probability estimates the average number of attractive interactions – sustaining protein aggregation – that each residue along the protein sequence establishes within bulk phase-separated droplets per protein replica. Thus, we can compare the maps of the same system for both the reference model and the ageing model (after reaching a plateau in the number of cross-$\beta$-sheet transitions over time) to explore the effect of accumulation of inter-peptide $\beta$-sheet nuclei inside matured condensates. From these probabilities, we can infer a binding free energy variation (if assuming that aged droplets at extremely long timescales can still be equilibrated):

$$\frac{\Delta G}{k_B T} \approx -ln\left(\frac{P_g}{P_l}\right), \tag{S15}$$

where $P_g$ and $P_l$ are the residue contact map probabilities for the ageing and reference models respectively, $k_B$ is the Boltzmann constant, and $T$ is the temperature. It is worth noting that the calculation using Eq. S15 can be flawed if any residue-residue contact never happens, either in the numerator or the denominator. To sort out such numerical drawback, we rescale both probabilities by adding one in both maps so the divergent points are reduced to a reasonable value. Additionally, when computing the difference in free energy for the A-LCD-hnRNPA1/polyU mixture (Fig. 5c of the main text), we artificially set the value of polyU-polyU interactions at a constant value of 1 for convenience. Since Uridine-Uridine interactions cannot enhance their binding due to structural transitions, and the contacts between polyU strands are almost negligible (but still non-zero) their consideration in the calculation is irrelevant (and would only add statistical noise without providing any physical insight).

### SUPPLEMENTARY SECTION VIII. CALCULATION OF TRANSPORT PROPERTIES WITHIN PHASE-SEPARATED CONDENSATES

To evaluate droplet viscosity and protein self-diffusion within bulk condensates, we run NVT simulations using a cubic box, and setting the system volume to the corresponding equilibrium bulk droplet density (further information on the simulation details is provided in Section SI D). We characterize the transport properties of pure condensates of A-LCD-hnRNPA1 and FUS-PLD at different temperatures using both the reference and the ageing models in Figs. 2a, 2b, 2c and 5d of main text. Additionally, the impact of increasing the polyU/A-LCD-hnRNPA1 mass ratio on the transport properties of aged condensates at T=0.97T$_c$ is also explored in Fig. 5d. Below, we provide the technical details of these calculations.

#### A. Estimation of viscosity through the shear relaxation modulus

From NVT simulations, we can compute both viscosity and protein diffusion in the condensate in separate ways. The shear viscosity can be straightforwardly calculated by integrating the relaxation modulus in time (see Chapter 7 of the book [38]):

$$\eta = \int_0^\infty dt\, G(t) \tag{S16}$$

In an isotropic system, we can compute the shear relaxation modulus $G(t)$ more accurately by using all the components of the pressure tensor ($\sigma_{\alpha\beta}$) as shown in Ref. [39]:

$$G(t) = \frac{V}{5k_BT}\left[\langle\sigma_{xy}(0)\sigma_{xy}(t)\rangle + \langle\sigma_{xz}(0)\sigma_{xz}(t)\rangle + \langle\sigma_{yz}(0)\sigma_{yz}(t)\rangle\right] + \frac{V}{30k_BT}\left[\langle N_{xy}(0)N_{xy}(t)\rangle + \langle N_{xz}(0)N_{xz}(t)\rangle + \langle N_{yz}(0)N_{yz}(t)\rangle\right], \tag{S17}$$

where $N_{\alpha\beta} = \sigma_{\alpha\alpha} - \sigma_{\beta\beta}$ is the first normal stress difference. This correlation can be easily computed by using the compute ave/correlate/long in the USER-MISC package of LAMMPS (version 21st of July of 2020) [3]. In all cases, the relaxation modulus presents an initial regime that mainly accounts for the intramolecular interactions, followed by a terminal region which corresponds to much slower relaxation modes, as those coming from intermolecular interactions and the relaxation of the large scale protein and RNA conformations. Due to the very wide range of timescales involved in the calculation and the noisy nature of the relaxation modulus in the terminal region obtained in the simulations, we follow a particular strategy to calculate our estimate of viscosity. At short times, $G(t)$ is smooth and the integral can be computed using numerical integration (trapezoidal rule). However, at longer times $G(t)$ presents more noise, and hence, we calculate the integral in that regime by first fitting $G(t)$ to a series of Maxwell modes ($G_i \exp(-t/\tau)$) equidistant in logarithmic time [40] and then by calculating the integral analytically. Our fit to the Maxwell modes is carried out with the help of the open-source RepTate software (version 1.1.1 20200602) [34]. Finally, viscosity is obtained by adding the two terms:

$$\eta = \eta(t_0) + \int_{t_0}^{\infty} dt\, G_M(t), \tag{S18}$$

where $\eta(t_0)$ corresponds to the computed term for short time-scales, $G_M(t)$ is the part evaluated via the Maxwell modes fit at long time-scales, and $t_0$ is the time that separates both (i.e., vertical dotted line in Fig 2b of the main text). The symbol size shown in Fig. 2d of the main text have been estimated from the error of the Maxwell mode fits to the value of $G(t)$ obtained in our simulations.

#### B. Calculation of the diffusion coefficient

The protein diffusion coefficient inside the condensates is obtained through the mean squared displacement (MSD) of the protein center of mass. After a subdiffusive regime (i.e., ∼1 molecular diameter), proteins exhibit a diffusive behavior and then the MSD of the center of mass can be measured via:

$$\left\langle (\mathbf{R}_{\mathrm{CM}}(t) - \mathbf{R}_{\mathrm{CM}}(0))^2 \right\rangle = 6D_c t, \tag{S19}$$

where $R_{CM}$ indicates the center of mass of a given protein at different times, and $D_c$ accounts for the diffusion coefficient. In order to get an accurate estimate of the MSD, the same correlator technique employed in the calculation of the relaxation modulus has been used [39]. By plotting the MSD divided by $6t$, the function shows a plateau at long timescales that provides the value of the diffusion coefficient of the proteins ($D_c$). We note that condensate viscosity and protein diffusion (shown in Figs. 2b, 2c, 2d and 5d of the main text) were evaluated through distinct trajectories which lasted between 3 to 4 $\mu$s.

### SUPPLEMENTARY SECTION IX. REFERENCES

### REFERENCES

[1] S. Das, Y.-H. Lin, R. M. Vernon, J. D. Forman-Kay, and H. S. Chan, Proceedings of the National Academy of Sciences **117**, 28795 (2020).
[2] G. L. Dignon, W. Zheng, Y. C. Kim, R. B. Best, and J. Mittal, PLoS computational biology **14**, e1005941 (2018).
[3] S. Plimpton, Journal of computational physics **117**, 1 (1995).
[4] R. M. Regy, G. L. Dignon, W. Zheng, Y. C. Kim, and J. Mittal, Nucleic Acids Research **48**, 12593 (2020).
[5] G. Krainer, T. J. Welsh, J. A. Joseph, J. R. Espinosa, S. Wittmann, E. de Csilléry, A. Sridhar, Z. Toprakcioglu, G. Gudiškytė, M. A. Czekalska, *et al.*, Nature Communications **12**, 1 (2021).
[6] H. S. Ashbaugh and H. W. Hatch, *Journal of the American Chemical Society*, Journal of the American Chemical Society **130**, 9536 (2008).
[7] L. H. Kapcha and P. J. Rossky, Journal of molecular biology **426**, 484 (2014).
[8] J. A. Joseph, A. Reinhardt, A. Aguirre, P. Y. Chew, K. O. Russell, J. R. Espinosa, A. Garaizar, and R. Collepardo-Guevara, Nature Computational Science **1**, 732 (2021).
[9] X. Wang, S. Ramírez-Hinestrosa, J. Dobnikar, and D. Frenkel, Physical Chemistry Chemical Physics **22**, 10624 (2020).
[10] P. Robustelli, S. Piana, and D. E. Shaw, Proceedings of the National Academy of Sciences **115**, E4758 (2018).
[11] H. J. Berendsen, D. van der Spoel, and R. van Drunen, Computer physics communications **91**, 43 (1995).
[12] M. P. Hughes, M. R. Sawaya, D. R. Boyer, L. Goldschmidt, J. A. Rodriguez, D. Cascio, L. Chong, T. Gonen, and D. S. Eisenberg, Science **359**, 698 (2018).
[13] W. Humphrey, A. Dalke, and K. Schulten, Journal of molecular graphics **14**, 33 (1996).
[14] B. Hess, Journal of chemical theory and computation **4**, 116 (2008).
[15] T. Darden, D. York, and L. Pedersen, The Journal of chemical physics **98**, 10089 (1993).
[16] S. Nosé, The Journal of chemical physics **81**, 511 (1984).
[17] M. Parrinello and A. Rahman, Journal of Applied physics **52**, 7182 (1981).
[18] J. S. Hub, B. L. De Groot, and D. Van Der Spoel, Journal of chemical theory and computation **6**, 3713 (2010).
[19] A. J. Ladd and L. V. Woodcock, Chemical Physics Letters **51**, 155 (1977).
[20] T. Schneider and E. Stoll, Physical Review B **17**, 1302 (1978).
[21] S. Plimpton, Journal of Computational Physics **117**, 1 (1995).
[22] S. Nosé, The Journal of Chemical Physics **81**, 511 (1984).
[23] W. G. Hoover, Phys. Rev. A **31**, 1695 (1985).
[24] H. Kamberaj, R. Low, and M. Neal, The Journal of chemical physics **122**, 224114 (2005).
[25] A. Stukowski, Modelling and simulation in materials science and engineering **18**, 015012 (2009).
[26] R. García Fernández, J. L. F. Abascal, and C. Vega, The Journal of Chemical Physics **124**, 144506 (2006).
[27] J. R. Espinosa, E. Sanz, C. Valeriani, and C. Vega, Journal of Chemical Physics **139** (2013), 10.1063/1.4823499.
[28] J. S. Rowlinson and B. Widom, *Molecular theory of capillarity* (Courier Corporation, 2013).
[29] J. A. Zollweg and G. W. Mulholland, The Journal of Chemical Physics **57**, 1021 (1972).
[30] A. Garaizar, J. R. Espinosa, J. A. Joseph, G. Krainer, Y. Shen, T. P. Knowles, and R. Collepardo-Guevara, Proceedings of the National Academy of Sciences **119**, e2119800119 (2022).
[31] J. R. Gissinger, B. D. Jensen, and K. E. Wise, Polymer **128**, 211 (2017).
[32] S. K. Sukumaran, G. S. Grest, K. Kremer, and R. Everaers, Journal of Polymer Science Part B: Polymer Physics **43**, 917 (2005).
[33] M. Doi, S. F. Edwards, and S. F. Edwards, *The theory of polymer dynamics*, Vol. 73 (oxford university press, 1988).
[34] V. A. Boudara, D. J. Read, and J. Ramírez, Journal of Rheology **64**, 709 (2020).
[35] F. Eisenhaber, P. Lijnzaad, P. Argos, C. Sander, and M. Scharf, Journal of computational chemistry **16**, 273 (1995).
[36] H. Wadell, The Journal of Geology **43**, 250 (1935).
[37] A. R. Tejedor, A. Garaizar, J. Ramírez, and J. R. Espinosa, Biophysical Journal **120**, 5169 (2021).
[38] M. Rubinstein, R. H. Colby, *et al.*, *Polymer physics*, Vol. 23 (Oxford university press New York, 2003).
[39] J. Ramírez, S. K. Sukumaran, B. Vorselaars, and A. E. Likhtman, The Journal of chemical physics **133**, 154103 (2010).
[40] A. E. Likhtman, Macromolecules **38**, 6128 (2005).
