## Supplementary material for "Protein structural transitions critically transform the network connectivity and viscoelasticity of RNA-binding protein condensates but RNA can prevent it": Description of Supplementary Movies

### Description of Additional Supplementary Files

**Supplementary Movie 1**:

Primitive path analysis applied to an aged FUS-PLD condensate at T/Tc,FUS = 0.785 revealing the topology of the underlying inter-protein -sheet binding network which has completely percolated through the phase-separated system. The aged condensate is kinetically arrested and presents gel-like behaviour.

**Supplementary Movie 2:**

Primitive path analysis applied to an aged FUS-PLD condensate at T/Tc,FUS = 0.861 revealing the topology of the underlying inter-protein -sheet binding network which has not fully percolated. The phase-separated condensate presents liquid-like behaviour at long timescales.
